# Reappraisal of the place of cultivated plants in the carbon budget

**DOI:** 10.1101/2025.05.17.654640

**Authors:** Arnaud Muller-Feuga

## Abstract

The impact of agriculture on the climate remains underestimated due to the systematic exclusion of annual crops from carbon budgets. Considered too ephemeral, these crops are nevertheless responsible for the absorption and storage of approximately one-third of the carbon biofixed by photosynthesis, with half-lives that are not limited to a single season but extend on average over 8.9 years. The kinetics of variation in carbon capture and release by cultivated plants over the half-century were simulated to complete the probabilistic calculation of the carbon budget components. In 2023, all cultivated plants (crops, grasslands, and forest plantations) had a stored carbon half-life of 17.6 years. They had removed 39.2+-0.5 billion tonnes of CO₂ (GtCO_2_) per year from the atmosphere, more than global emissions from hydrocarbon combustion. The time distribution allowed by this simulation suggests that cultivated plants would have biofixed a cumulative net total of 455 GtCO_2_ in 2023, or 14% of the mass of atmospheric CO_2_. Given the importance of this anthropogenic carbon capture and storage by cultivated plants, both in duration and quantity, rural activities should be integrated into carbon budgets and the resulting climate strategies, and recognized as a carbon capture and storage (CCS) device in carbon cycle regulation policies. This recognition would allow for the fair valuation of the work of farmers and foresters as part of the ecological transition, particularly through remuneration mechanisms such as carbon credits.

## Introduction

The evolution of Earth’s climate has long been a subject of particular attention. Decision-makers strive to identify the most influential factors in order to predict their evolution and act on them, when possible, particularly if they pose risks to humanity. The Intergovernmental Panel on Climate Change (IPCC) was established in 1988 by the World Meteorological Organization (WMO) and the United Nations Environment Programme (UNEP). Its mandate is to impartially assess international scientific, technical, and socio-economic information on climate change. Since its first report in 1990, and regularly every 5-6 years, the IPCC issues a synthesis report that summarizes the contributions of three working groups involving hundreds of specialists. These assessment reports serve as a reference for scientists and decision-makers around the world and are the basis for policies implemented by most OECD and European Union countries.

This scientific production, at the heart of international climate negotiations, aims to alert decision-makers and civil society. Since the IPCC’s creation, all of its reports have been adopted in plenary session by all 192 countries represented. The IPCC’s annual budget of approximately five million euros in 2012 increased to six million euros in 2021. It is funded by the 195 UN member states who contribute “independently and voluntarily.” Several hundred scientists participate in the preparation of reports that present the situation of climate change as the consequence of emissions, particularly of carbon dioxide, and recommend binding restrictions on the use of fossil fuels.

Global warming, which has been occurring for half a century at a rate of 0.19°C every 10 years (Met Office Hadley Centre, 2024), is believed to be a greenhouse-type process and the consequence of the emission of CO_2_ produced by the combustion of fossil fuels into the atmosphere since the beginning of the industrial era. The atmospheric CO_2_ content (ACC) has increased from 270 parts per million by volume (ppm) in the mid-19th century to 420 ppm. This increase appears to be due to the release of products from the combustion of fossil fuels, coal, oil, and gas, which has intensified since then. The increase in global temperatures observed over the same period is believed to be due to these emissions, which are presented as pollution.

According to the most recent scenarios, which assume a warming of 1.5°C, global sea levels could rise by between 0.26 and 0.77 m by the end of this century. Such an increase would be detrimental to populations living on low-lying coasts. Strong measures should be taken quickly to prevent the enrichment of the atmosphere with carbon dioxide from causing lasting global climate change in the coming decades. According to the 5th report, a doubling of the ACC would cause a 2.8°C increase in the average tropospheric temperature, currently 14.5°C. This average warming is significant compared to that following a glaciation, which is 4°C.

Most governments have embraced this view and renew their endorsement at the annual COP (Conference of the Parties). One consequence is the allocation of tradable emission permits that must not be exceeded subject to penalty (Kyoto Protocol - 1997) within the framework of a global carbon market. In order to move closer to neutrality, emitting activities are encouraged to purchase such permits from those that carry out carbon capture and storage (CCS) emitted, avoided, or removed directly from the atmosphere. Sites emitting more than 0.1 MtCO_2_/year must offset their emissions with a CCS system or purchase emission permits or carbon credits.

CCS through geological burial for several centuries is being adopted by most industrialized countries to achieve carbon neutrality by 2050. It appears to be the means to fulfill the promises made by governments and renewed at successive COPs. If the agenda is not compliant and the promises are not kept, penalties are imposed. However, countries sometimes denounce previously signed binding agreements, as was the case with Canada, which denounced the Kyoto Protocol in 2011.

The development of CCS through landfilling is sometimes perceived as compatible with the continued use of fossil fuels, raising questions about its actual contribution to carbon neutrality. Its principle is also widely criticized by those who would like to “decarbonize” the global economy and who view CCS and emission permit quotas as pollution rights.

Carbon capture and storage in plant products marketed by crops, livestock, and forestry, based on FAO (n.d.) statistics, has been estimated (Muller-Feuga, 2024a) at 21 billion tonnes of CO_2_ (GtCO_2_ or PgCO_2_). Subsequently, the non-commercial parts carrying these productions were taken into account (Muller-Feuga, 2024b), which increased the carbon capture and storage by whole cultivated plants (CCSP) to 41.0±0.6 GtCO_2_/year and the average storage duration weighted by dry weight to 26.3±2.0 years in 2022. These figures were much higher than those in the literature: 3.2 times the continental sink of Friedlingstein et al. (2022) estimated at 12.8±3.3 GtCO_2_/year, and 2.1 times that of Pan et al. (2024) estimated at 19.8 GtCO_2_/year. The analysis was completed by that of the evolution over 50 years (Muller-Feuga, 2025) of the CSCP and the restitutions by respiration and combustion to which they give rise. It showed that plant cultivation constituted an average sink of 39.9 GtCO_2_/year removed from the atmosphere during the ten years preceding 2022 and offset the 36 GtCO_2_ emitted by the combustion of fossil hydrocarbons. It also showed that the global ocean was a growing source of 10.6 GtCO_2_/year on average during the last decade.

These differences appear to be due to the failure to take annual plant crops into account in carbon budgets. We noted a specific provision in the document describing the methods and guidelines (IPCC, 2006a and 2006b) for researchers. In Chapter 5: “Cropland,” is written “*The change in biomass is only estimated for perennial woody crops. For annual crops, increase in biomass stocks in a single year is assumed equal to biomass losses from harvest and mortality in that same year - thus there is no net accumulation of biomass carbon stocks”*.

This is confirmed by the 2019 edition of the guidelines (IPCC, 2019). Among plants, it is explicitly recommended to consider only perennial woody crops such as tea, coffee, oil palm, coconut, rubber trees, fruits and nuts, and polycultures such as agroforestry systems (e.g. Walker et al., 2021). This significantly reduces the proportion of plants entering the carbon budgets at the origin of the climate protection policy. However, carbon remains included in plant biomass during the few months to a few centuries that separate the start of growth from mineralization by digestion or combustion causing its return to the atmosphere in the form of CO_2_.

Cereals, including corn, wheat, and rice, which account for just under half of global agricultural production, have virtually infinite storage life under the right humidity and temperature conditions, whether as grains or processed into biscuits, pasta or noodles. In addition to stabilizing prices, this allows for their long-distance transportation and the creation of strategic reserves for populations affected by crises or located far from production areas.

According to the FAO Cereal Supply and Demand Bulletin (FAO, 2025), global production forecasts were 2,849 million tonnes in 2024, capturing 3.8 GtCO_2_ from the atmosphere.

Utilization forecasts for 2024-2025 are projected at 2,868 million tonnes. By the end of the 2025 season, global cereal stocks are expected to reach 873.3 million tonnes. The stock-to-use ratio of 30.1% suggests a rotation period of approximately three years.

For rice, the main Asian cereal, the closing stock of each production year represents one-third of utilization, which helps control world prices (FAO, 2018). This assumes a complete rotation at least every three years. Japan recently opened its strategic rice reserves to cope with soaring prices, which nearly doubled following the poor 2023 harvest. Of a total of approximately one million tons, 210,000 tons have been sold (The Mainichi, 2025). This is the first time since their creation in 1995 that these reserves have been depleted, representing a storage period of 30 years.

More generally, the global rice consumer price index shows a persistent upward trend, which is encouraging countries to import and build up reserves. The example of the Sahel demonstrates the extent to which food reserves are an important component of food security. The number of undernourished people in the world was 733 million in 2023, representing 9.1% of the global population (FAO, 2024). The rotation of global food reserves is a strategic issue that requires special attention to ensure food security in the face of current and future crises.

Thanks to their intrinsic capacities and conservation techniques, stocks constituted by annual plants persist well beyond the harvest year, and there is no justification for excluding them from carbon balances given the importance of their contribution. Cultivated plants fix carbon dioxide from the atmosphere and constitute a CCS device both in terms of their quantities and their duration. Here we provide the necessary updates and corrections and examine their consequences.

## Material and methods

### 1. Photosynthesis

Plants directly capture atmospheric CO_2_, then polymerize the carbon into biomass using the photosynthesis process that gave rise to life on Earth 2.5 billion years ago. The resulting carbon feeds heterotrophs, which degrade it through respiration. It is also mineralized through ignition and combustion. These polymerization and mineralization reactions are grouped under the same reversible chemical equation (1).

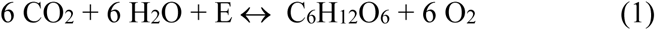

E is the visible light energy in the direction of photosynthesis (from left to right) and the metabolic or combustion energy in the direction of respiration (from right to left). This equation expresses that CO_2_ consumption, oxygen production, and the production of organic matter in the form of hexoses correspond molecule for molecule.

Reaction (1) implements a chain of enzymatic processes involving carbonic anhydrase and rubisco for the entry of this gas into the plant cell, followed by a series of reactions modulated by visible light. This equation lacks other essential but minor assimilable elements such as nitrogen, phosphorus, iron, etc. However, this equation is retained here because we limit our analysis to carbon.

Hexose is the basic building block of plant matter, and its quantity can be measured by the dry plant biomass (dpb) according to the stoichiometry of the chemical reaction (1). The carbon/dry-matter ratio (C/dm) it provides actually varies with the type of biomass and should be adjusted for each case. Indeed, it is 0.40 for glucose, 0.44 for cellulose, 0.64 for hemicellulose, and 0.66 for lignin. Despite this variation and for the sake of conservative simplification, we retain the value of 0.4 provided by equation (1) for all biomasses. One ton of dry plant biomass contains 0.4 tons of carbon (C/dm) and required 1.47 tons of CO_2_ (CO_2_/dm) taken from the atmosphere. In the opposite direction of the reaction, one ton of dry organic matter releases 1.47 tons of CO_2_ during its mineral degradation through respiration or combustion.

Although the efficiency of converting solar energy into biomass is in the order of a few percent, the resulting plant production is a significant carbon sink. Controlling all or part of the food chain, agriculture, livestock farming, forestry, hunting, fishing, and aquaculture feed, clothe, warm, shelter, entertain, etc… humanity. Here, we consider the net primary production, after autotrophic respiration, of these activities, which remove carbon dioxide from the atmosphere and store it in the form of biomass. As such, they are involved in the carbon cycle as providers of food or fuel.

A distinction must be made between stock and flow, the latter being the derivative of the former’s change over time. In annual budgets, only the flow is considered to measure atmospheric CO_2_ fixation. The flow is proportional to the quantity of plant biomass produced annually and then harvested for later use elsewhere. It is this flow that creates a CCS system that must be considered in the same way as those by burying (Muller-Feuga, 2024a).

### 2. The atmospheric CO_2_ budget

The results of the calculation of CO_2_ world absorption (AW) by plants and CO_2_ world emission (EW) by those that feed on them are compared with the other components of the global atmospheric carbon budget. They are the variation in atmospheric CO_2_ (ACV) and emissions from fossil fuel combustion (EFOS) that are fully documented in public statistics and allow us to specify the CO_2_ exchanges between continents, the atmosphere, and the ocean. Given the extreme complexity of the latter’s behaviour, heterogeneous in three dimensions and over time, the oceanic CO_2_ contribution OC is the unknown of equation (2), where the sources are positive and the sinks negative.

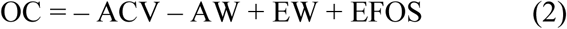

The following assessment of EW and AW is of an accounting nature and does not use numerical models based on satellite or field-measured data.

### 3. Quantities captured

Emerged land is cultivated to meet the needs of humanity and its livestock for food plants (cereals, vegetables, fruits, etc.), textiles (cotton, flax, hemp, etc.), ornamental plants (tobacco, grapes, flowers, etc.), heating, and construction. In the following, we use FAO statistics (n.d.) relating to agricultural, livestock, forestry, aquaculture, and fisheries production. The carbon material thus obtained feeds animals and microorganisms that mineralize it through digestion and respiration. Some of it is directly mineralized through deliberate or accidental combustion. These productions, responsible for continental CO_2_ absorptions by AW photosynthesis, are described in Muller-Feuga (2024b) and updated here in terms of captured quantities and storage durations.

The quantity CS of CO_2_ captured and stored by a set n of plant biomasses of fresh weight Pi, is equal to the sum ∑ of the anhydrous weights of the harvests multiplied by the CO_2_/dm ratio, i.e. 1.47 noted k, in accordance with equation (3), where TE_i_ is the water content of the biomass P_i_.

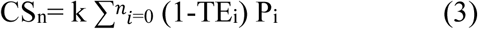

### 4. Storage duration

Freshly harvested fruits and vegetables have a reduced transport and marketing time in order to preserve their taste and freshness. Temperature control during transport and storage can significantly increase the time between harvest and consumption. This shelf life varies from 3 days to a few weeks at room temperature or in the refrigerator. It can go up to 3 months, which allows to get through the winter. In the freezer, food generally keeps for a year or more. In addition to temperature, the shelf life of food depends on its water content, which promotes its decomposition. This content varies: 10 to 20% in cereals, 60 to 75% in meats and animal flesh, 80 to 90% in fresh fruits and vegetables. Reducing the water content to extend storage time can be achieved through various techniques such as drying, brining, and vacuum packaging.

The probabilistic simulation of the distribution over time of plant carbon (Muller-Feuga (2024b) is taken up here. The history of carbon in agricultural and forestry products is divided into the period of capture by photosynthesis (CP) and the period of restitution by mineralization (RM). Harvest marks the transition between the two periods CP and RM. The first period (CP), which separates the beginning of plant growth, sowing, planting, or previous harvest (n-1), from harvest (n), gradually builds up the carbon pool through photosynthetic capture of CO_2_ from the atmosphere. This period lasts from a few months for annual plants to a few decades for trees. Since the weight growth over time of fruits and vegetables generally exhibits an S-shaped curve (e.g., Tijero et al., 2021), the amount of carbon captured and stored is assumed to be distributed over time according to an increasing normal distribution.

During the second period (RM), mineralization occurs after harvest, during which the polymerized carbon is mineralized and returned to the atmosphere in the form of CO_2_ through food ingestion followed by digestion and respiration, fermentation, or combustion. The organic matter is stored on shelves, in bulk, or in the soil, then ingested, fermented, or burned. This period varies from a few days for perishable fruits and vegetables to several centuries for structural timber. We assume that this CO_2_ emission is distributed over time according to a decreasing normal distribution.

Since the normal distributions are centered, the mean and median carbon storage half-lives (DSC) are equal to the sum of half the maximum growth time (PC/2) and half the maximum restitution time (RM/2) according to formula (4).

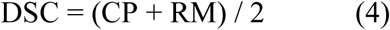

### 5. Simulation of CCSP kinetics

The normal distribution equations simulating the kinetics of capture and restitution of atmospheric CO_2_ are, for capture :

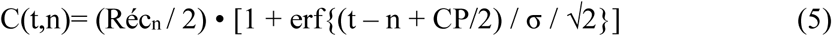

and for restitution :

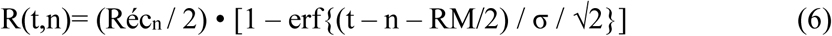

where t is the historical time in years, n is the harvest year, Rec_n_ is the atmospheric CO_2_ stock captured during the harvest in year n, erf is the error function of the centered normal distribution, and σ is the curvature of the curve. If t ≤ n, the CP capture duration applies and the erf function is added. If t > n, the RM restitution duration applies and the erf function is subtracted.

This probabilistic simulation (see Figure 2) is applicable to plants with a single harvest, such as annual plants and trees felled for fuel and construction wood, and to perennial plants, such as sugarcane, tea, coffee, cocoa, oil palm, rubber trees, fresh and dried fruits, which are exploited by multiple annual harvests. In the latter case, the CP capture period covers previous multiple harvests, which varies depending on the species between 2 and 80 years.

Let us remember that perennial woody crops are the only ones considered in most carbon budgets, excluding annual plants.

### 6. Other calculation conditions

To calculate storage durations and captured CO_2_ quantities, assumptions are made regarding water content, protein content, and the durations of the CP and RM periods, based on the most relevant data from the literature. The quantities of captured and restituted CO_2_ are calculated using formula (3) by deducting the water content TE from the fresh weight provided by the statistics. The CP and RM durations are assigned to each biomass, then weighted by the anhydrous weights for each of the three biomass groups: crops, fodder, and forestry. The dispersion of the results is expressed by their standard deviation.

The water contents TE_i_ documented by various cross-referenced sources allow the dry weight P_i_ of these plant products to be calculated using formula (3). Figure 1 shows the variation of the maximum restitution duration RM of crop products placed on the market as a function of their water content TE. We distinguish between fresh fruits and vegetables for which the RM is from a few days to a few months, semi-preserved products for which the RM is from a few months to a few years, and dried products for which the RM is from a few years to indefinitely.

**Figure 1:**
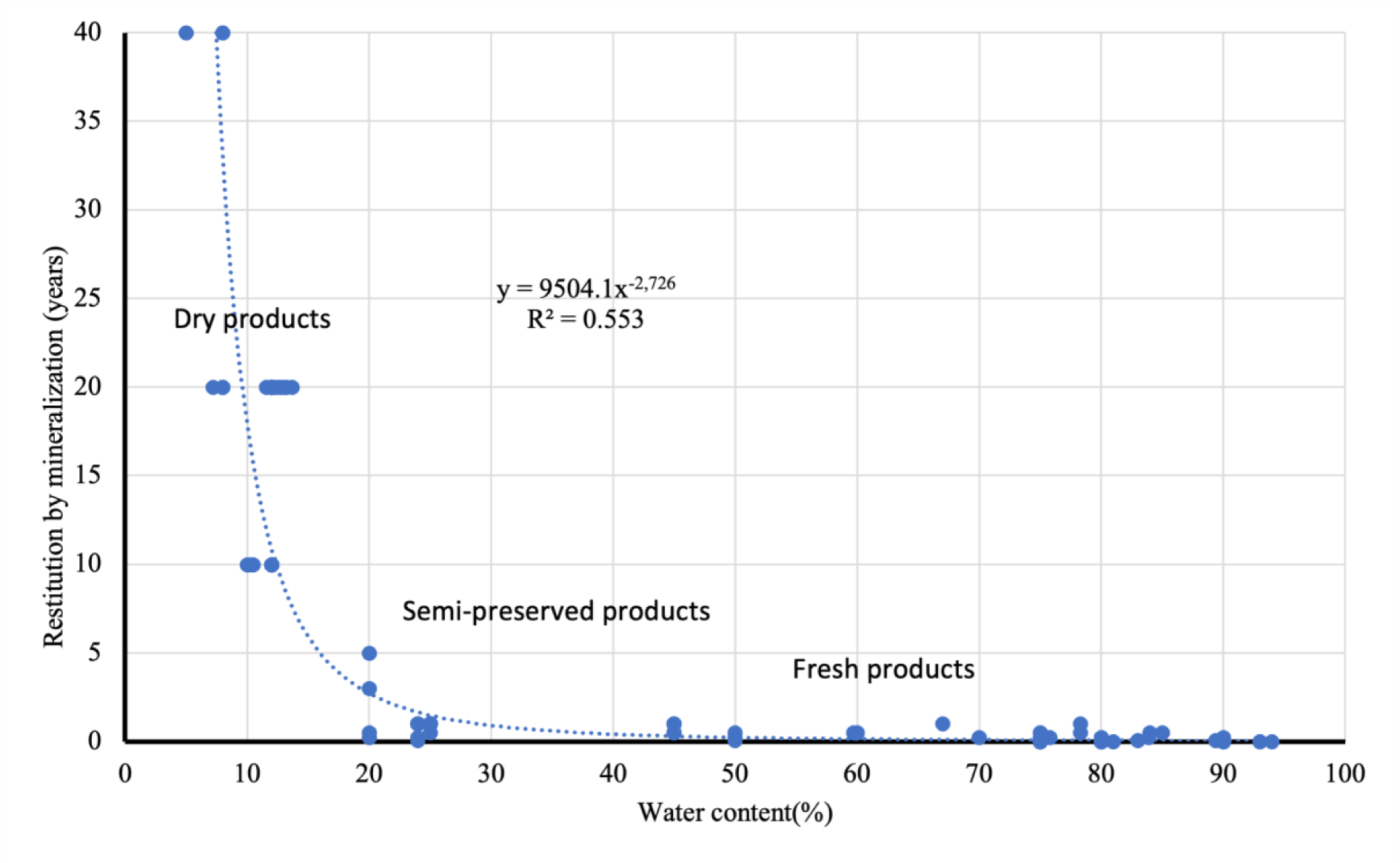
Post-harvest restitution times (RM) of commercial crop products vs their water content in 2023.

Since the nutritional value and safety (freshness) of foods decrease with age, consumers are guided in their choices to ensure food safety and limit waste. Best-before dates (BBD) are mandatory information on food products. We considered our maximum restitution duration RM to be equal to these BBDs when they exist.

The primarily food use of agricultural products prolongs carbon storage in the form of animal biomass and their waste, whose carbon contents are approximately 18 and 9% by weight, respectively. The lifespan of animal carbon before being released to the atmosphere through respiration ranges from one week in the liver to six weeks in hair (Tieszen et al., 1983). Fecal matter in the form of compost or manure is mineralized in the environment, with a few months of estimated carbon lifetime. This animal extension is not taken into account here, which minimizes the carbon storage duration.

The average carbon lifetimes of necromasses correspond to the average carbon residence time before mineralization by soil decomposers. For forests, these retention times vary between 0.9 and 152 years (Wang et al., 2017). They increase with latitude (x4) and decrease with temperature (x10) and rainfall (x4). We selected the durations for temperate regions, which represent a compromise between high and low latitudes, where the maximum organic carbon residence time is 75 years for logged forest soils and 40 years for crop and grassland soils (Balesdent and Recous, 1997; Pellerin et al., 2019).

### 7. Matrix calculation

The theoretical kinetic of annual CCSPs by whole plants modeled by equations (5) and (6) allows us to construct three 118x127 matrices, which we call [C(t,n)] for capture, [R(t,n)] for restitution, and [C(t,n) + R(t,n)] for the complete cycle, where t is between years 1961 and 2079, and n is between years 1932 and 2059. Rows t contain the stocks formed by successive harvests as a function of time n, and columns n contain the captured, harvested, and then restituted stocks of year n as a function of time t (see Figure 2).

**Figure 2.**
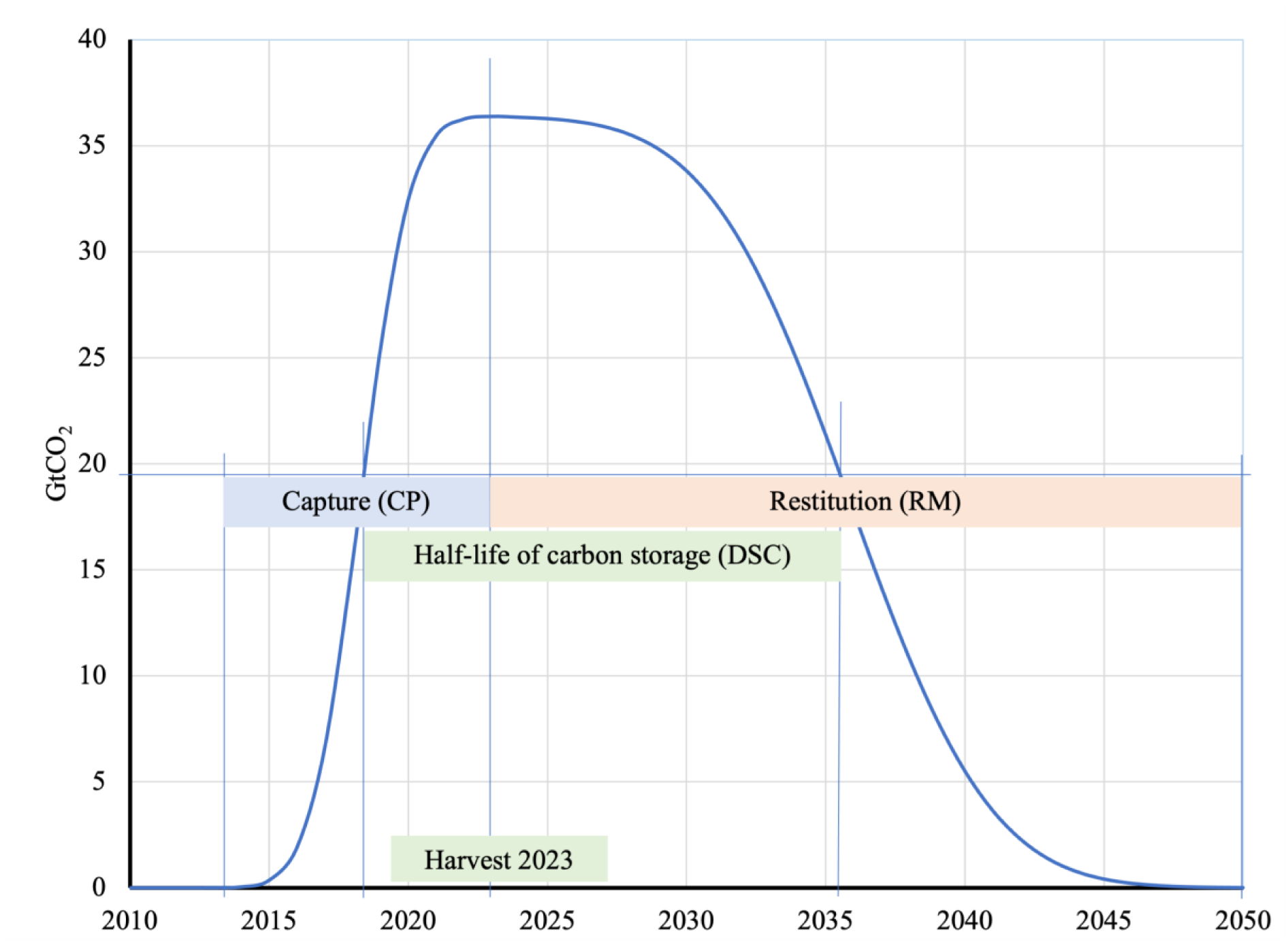
: Simulated kinetics of carbon stock variation (GtCO_2_) in whole plants bearing the 2023 harvests during capture by photosynthesis (CP) and restitution by mineralization (RM), and half-life of the carbon stock (DSC).

The continental balance excluding fossil emissions is equal to – AW + EW, where AW is the flux captured by plants and EW the flux restituted from plants. Numerically, it is taken equal to the variation from one year to the next, noted Δ, of the sum of the stocks of line t of the matrix [C(t,n) + R(t,n)], noted ∑_t_, reduced by the sum of the stocks of line t-1 of the same matrix. Formula (7) makes it possible to calculate the EW_t_ restitutions.

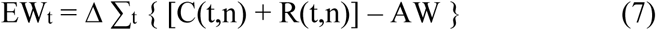

The quantity AW_t_ of CO_2_ captured in year t is equal to the sum of the harvests of year t, noted Réct, and the intermediate captures of the plants growing during the year and preparing the harvests to come, noted Cin_t_, in accordance with formula (8).

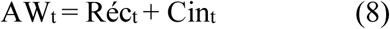

The intermediate captures Cin_t_ of year t are equal to the variation from one year to the next, noted Δ, of the sum of the captures of line t of [C(t,n)] after deduction of the harvests Réc_t_, noted ∑_t_, reduced by the same sum of line t-1, i.e. expression (9).

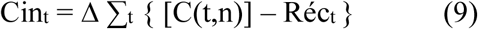

The amount of CO_2_ taken from the atmosphere by cultivated plants in year t is equal to the sum of row t of the matrix [C(t,n) + R(t,n)] for n ≤ t.

## Results

The total amount of CO_2_ captured by whole plants is calculated based on FAO decadal statistics (n.d.) describing marketed agricultural, livestock, and forestry products. These quantities are converted to anhydrous products, then multiplied by their carbon content (40%), then by the CO_2_/C mass ratio (3.37), then by the whole plant/commercial part ratio.

### 1. Crops 2023

Production statistics for the 160 global crop products include herbaceous and shrubby plants grown for food (cereals, vegetables, fruits, etc.), textile fibers (cotton, flax, hemp, etc.), or pleasure (tobacco, grapes, flowers, etc.). Fresh weights are described under “Production,” then “Crops and Animal Products”. Table 1 shows the durations and quantities of these commercial products broken down into the three categories described above. In 2023, the average dry weight-weighted water content (TE) of crop products was 45.6%. We considered the maximum restitution duration RM for cereals and other dry products to be 20 years.

**Table 1:** CP capture durations, RM restitution durations, dry weights, and CCSPs of the 160 global crop products marketed in 2023 (source: FAO, n.d.).

|  | CP<br>year | RM<br>year | Dry weight<br>Mt | CCSP<br>GtCO<sub>2</sub>/year |
| --- | --- | --- | --- | --- |
| Dry products | 9.3 | 17.7 | 3,926.3 | 5.76 |
| Semi-preserved products | 6.9 | 0.8 | 421.2 | 0.62 |
| Fresh products | 7.8 | 0.1 | 1,030.2 | 1.51 |
| Total | 4.4 | 15.6 | 5,377.8 | 7.89 |

### 2. Fodder 2023

Fodders are mostly annual plants, the cuttings of which are stored for 0.8 to 3 years to wait for maturity and to get through unproductive periods. They are transformed into meat, offal, eggs, honey, etc. by the animals that feed on them and mobilize their carbon. The restitution in the form of CO_2_ is done by respiration of livestocks as well as by mineralization of their excrement and non-food products (skins, wax, silk). It lasts between 40 days (eggs) and 100 years (beeswax). The 48 global livestock products are listed in the form of quantities of meat, milk, eggs consumable by humans expressed in tons described in the statistics “Production - Quantity” and “Primary Livestock”. These products are mainly intended to supplement human diet with quality proteins and a set of micro-nutrients such as vitamins A, B-12, riboflavin and mineral salts. They result from the conversion of plants from grasslands, foliage, and crops by ruminant and monogastric animals. Protein levels vary between 3.2% for milk and 32% for meat products.

According to Mottet et al., 2017, global livestock consumed 6 Gt/year of anhydrous fodder in 2010. After updating based on demographics, these anhydrous fodders are calculated by applying an average conversion rate (dry weight of fodder/fresh weight of livestock product) of 4.33 for all species combined. Some of the crops described in the previous section are intended for animal feed. These are cereals (wheat, corn, sorghum, etc.) used in livestock feed. To avoid counting them twice, a 14% reduction is applied to fodder calculated on the basis of animal products marketed (Table 2). This is the share of fodder that can be consumed by humans, according to the work cited above.

**Table 2:** CP capture duration, RM restitution duration, anhydrous weights and CCSPs by the 48 global livestock products marketed in 2023 (source FAO, n.d.).

|  | CP<br>year | RM<br>year | Dry weight<br>Mt | CCSP<br>GtCO<sub>2</sub>/year |
| --- | --- | --- | --- | --- |
| Dry products | 0.9 | 18.5 | 145.4 | 0.21 |
| Semi-preserved products | 0.7 | 0.6 | 4,200.6 | 6.16 |
| Fresh products | 2.0 | 0.2 | 2,125.0 | 3.12 |
| Total | 1.3 | 0.8 | 6,461.5 | 9.48 |

### 3. Forestry 2023

Statistics on the quantities of wood produced worldwide are available in the “Forests” section of the FAO “Data” (n.d.). These “Production - Quantity” of firewood, sawlogs, pulpwood, and industrial wood, expressed in m^3^, are converted into dry tons (Table 3). For this purpose, a density of 0.6 t/m^3^ is used for conifers and 0.7 for non-conifers.

**Table 3:** CP capture duration, RM restitution duration, anhydrous weights and CCSPs by the 8 global forestry products marketed in 2023 (source FAO, n.d.).

|  | CP<br>year | RM<br>year | Dry weight<br>Mt | CCSP<br>GtCO<sub>2</sub>/year |
| --- | --- | --- | --- | --- |
| Dry products | 40.0 | 53.3 | 1,240.0 | 1.82 |
| Semi-preserved products | 40.0 | 2.0 | 4,200.6 | 6.16 |
| Fresh products | 0.0 | 0.0 | 2,125.0 | 3.12 |
| Total | 44.0 | 18.3 | 2,589.8 | 3.80 |

### 4. Other 2023

In addition to these products from cultivated plants, we should add the uptake by wild plants and animals. The stock of unexploited organic matter (primary forests, necromasses, soil heterotrophs, peats, hedges, etc.) is relatively stable, and its flow occurs at the margins of the carbon polymerization-mineralization cycle. Unexploited plant covers have a photosynthesis-respiration balance close to neutral over time, with what is photosynthesized during the day being degraded at night or in the shade of the canopy by decomposing organisms or by combustion. We did not take these carbon stocks into account, which gives our figures a conservative value. Similarly, we considered the variations in carbon stocks from aquatic products (fishing and aquaculture) and land-based animal populations (humans and livestock). They were lower by a factor of 1000 than those of the three main terrestrial plant productions, so they were subsequently neglected.

### 5. Non-commercial Parts

Non-commercial plant biomass inherent to crops, such as leaves and stems for the aboveground portion, and roots and exudates for the belowground portion, must also be taken into account. These parts remain in situ and enrich the soil with organic carbon. The storage half-lives DSC of these necromasses correspond to the average residence time of carbon before mineralization by decomposer organisms.

The weights of the aboveground parts were calculated based on an average harvest index of 0.42 (Hay, 1995). The root system is estimated to account for 30% of crop plant biomass, 65% of grassland plants, and 15% of total tree biomass (Blume et al., 2015). Consequently, the whole plant/commercial part ratios are 1.72, 2.07, and 1.57 for crops, forages, and forest products, respectively.

### 6. Storage durations 2023

Storage durations were calculated on these bases, weighted by the anhydrous biomasses concerned for the year 2023 (Table 4).

**Table 4:** Photosynthetic capture (PC) and mineralization restitution (MR) durations by commercial parts, non-commercial parts, and whole plants harvested in 2023, as means weighted by the anhydrous weights of the biomasses.

|  | Commercial parts |  | Non-commercial parts |  | Whole plants |  |
| --- | --- | --- | --- | --- | --- | --- |
|  | CP<br>(year) | RM<br>(year) | CP<br>(year) | RM<br>(year) | CP<br>(year) | RM<br>(year) |
| Crops | 4.4 | 15.6 | 4.4 | 40.0 | 4.4 | 25.8 |
| Forages | 1.3 | 0.8 | 1.3 | 40.0 | 1.3 | 21.1 |
| Forests | 44.0 | 18.3 | 44.0 | 75.0 | 44.0 | 38.9 |
| Mean | 10.7 | 10.1 | 10.7 | 44.6 | 9.4 | 25.8 |

The half-life DSC, according to formula (4), for carbon stocks constituted by all cultivated plants is 17.6 years. The DSC of exploited trees is the longest, at 41.4 years. The DSC of forage plants is the shortest, at 11.2 years, which is sufficient to be included in carbon budgets. Crops have an intermediate DSC of 15.1 years. Following these results, the theoretical kinetic of 2023 CCSP is simulated using equations (5) and (6) with the parameters in Table 5.

**Table 5:**
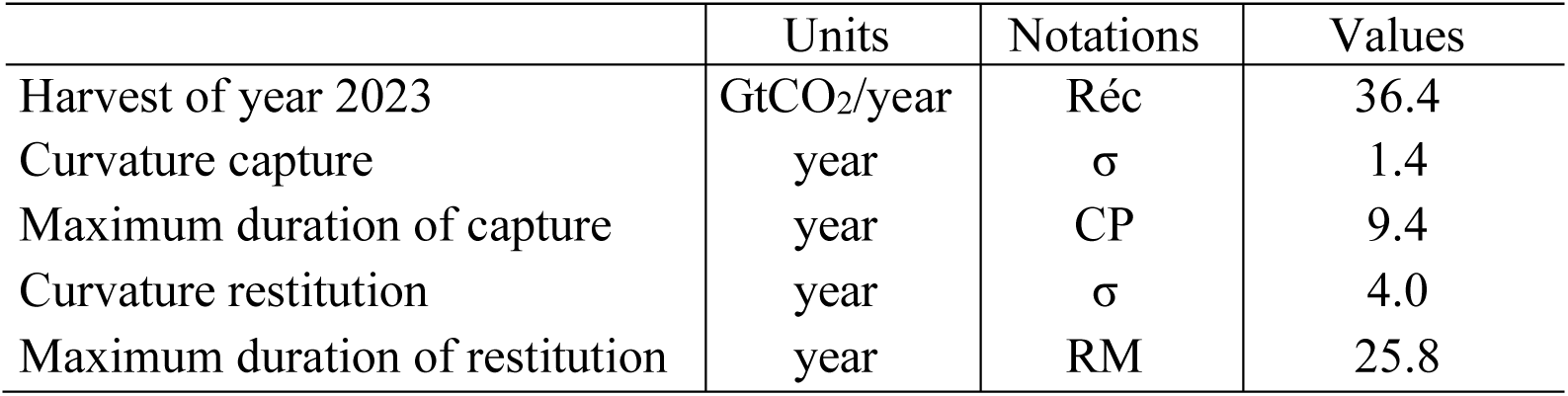
Parameters of the simulation using formulas (5) and (6) of the variation of carbon stocks as a function of time in 2023.

|  | Units | Notations | Values |
| --- | --- | --- | --- |
| Harvest of year 2023 | GtCO<sub>2</sub>/year | Réc | 36.4 |
| Curvature capture | year | $\sigma$ | 1.4 |
| Maximum duration of capture | year | CP | 9.4 |
| Curvature restitution | year | $\sigma$ | 4.0 |
| Maximum duration of restitution | year | RM | 25.8 |

Figure 2 shows the kinetics of variation in atmospheric CO_2_ captures and releases following the 2023 harvests. It describes the elements of the 2023 column vector of the matrix [C(t,n) + R(t,n)]. The CO_2_ stock removed by the 2023 harvest extends over approximately 40 years. The kinetics are highly asymmetric, with restitution occurring slower than capture.

This delay in restitution results in an accumulation of carbon stocks from successive harvests, the production of which is increasing.

### 7. Net balance 2023

In addition to the carbon stock of harvested plants, there is that of future harvests currently being built up. This intermediate capture, denoted Cin, was -2.7 GtCO_2_/year according to (9), bringing the total capture of global whole cultivated plants (AW) to -39.2±0.5 GtCO_2_ in 2023 (Table 6).

**Table 6:** Total quantities of CO_2_ captured and stored by the 2023 harvests in commercial products and whole plants of global agriculture, livestock, and forestry and their standard deviation (SD), in GtCO_2_/year.

| GtCO<sub>2</sub>/year | Commercial products | SD | Whole plants | SD |
| --- | --- | --- | --- | --- |
| Crops | 7.9 | 0.2 | 13.6 | 0.5 |
| Forages | 8.2 | 0.3 | 16.9 | 0.5 |
| Forests | 3.8 | 0.4 | 6.0 | 0.5 |
| Intermediate captures ($C_{in}$) | | | 2.7 | 0.8 |
| Total | 19.8 | 0.3 | 39.2 | 0.5 |

Formula (7) gives an EW restitution of 8.2±1.8 GtCO_2_/year in 2023, i.e. a net balance of global cultivated plants of -31.0±1.9 GtCO_2_/year taken from the atmosphere.

### 8. World CCSP over the half-century

Except for harvests Réc, the other parameters σ, CP, and RM in Table 5 were retained to simulate stock variations from 1932 to 2059. To describe the variations in global CCSPs over the past half-century (Figure 3), it was necessary to go back to 1940 to include all stocks in process of restitution in 1970, and to anticipate to 2030 to include all stocks in process of capture in 2023. To do this, we extrapolated to 1940 using the linear regression AW = 0.4198•n - 809.39 and to 2030 using the regression AW = 0.4883•n - 945.42, where n is the year considered. This is an acceptable hypothesis given the high value of the coefficients of determination (R^2^ ≥ 0.985).

**Figure 3.**
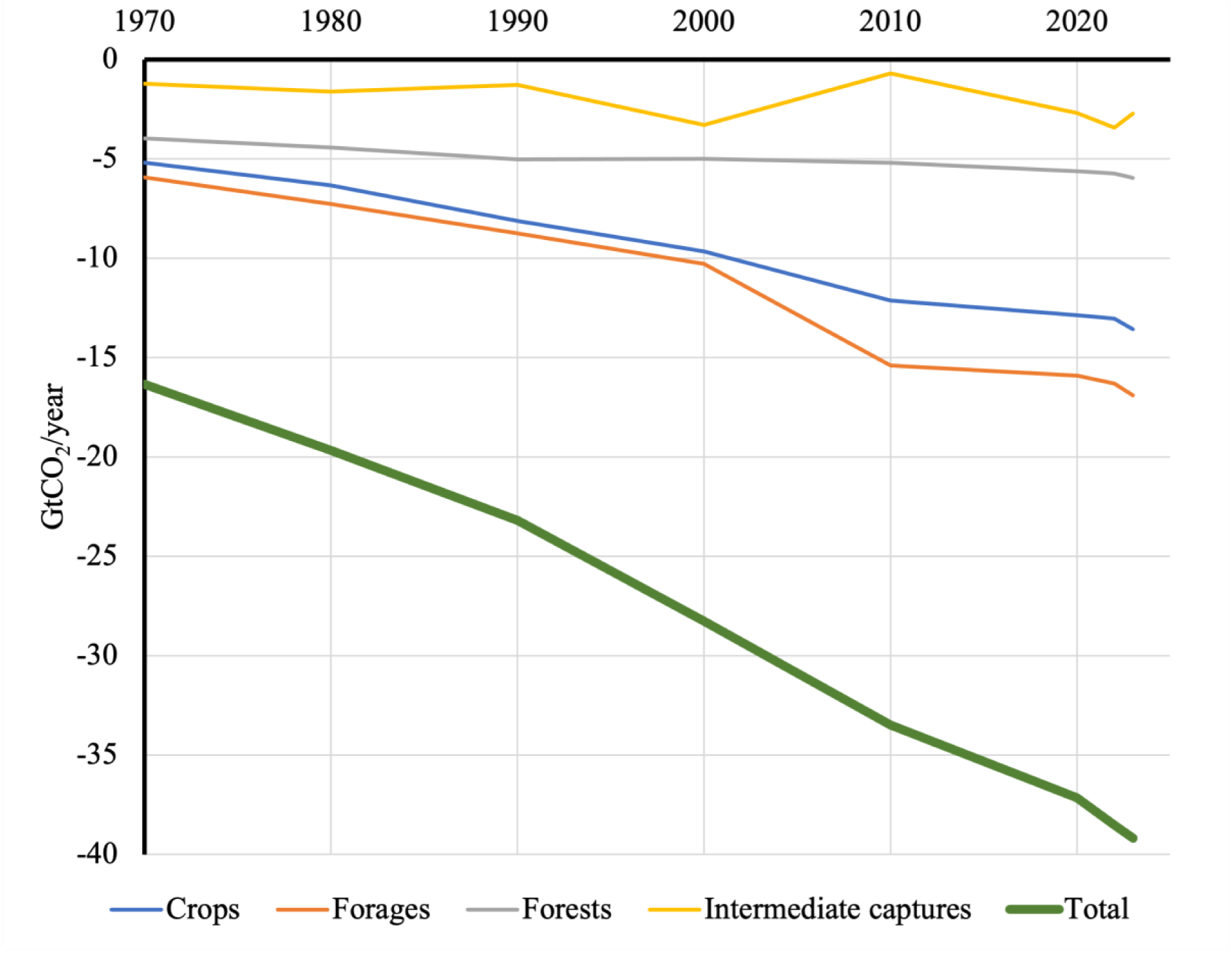
: Global CO_2_ uptake by whole plants cultivated from 1970 to 2023 (GtCO_2_/year; based on FAO, n.d.).

Recall that these CCSPs are sinks and that their values are negative in formula (2) of the atmospheric budget. Only an unexplained bump due mainly to fodder breaks the linearity around 2010. The final downward bracket marks the recovery following the COVID-19 pandemic.

The description of the kinetics of CCSPs of 2023 was extended (Figure 4) to the half-century 1970-2023 by assuming that only harvests Réc vary and that the other parameters in Table 5 remain unchanged for the years before and after 2023. This provision appears acceptable given that the relative proportions of agricultural and forestry products have not changed much over the half-century, unlike the quantities.

**Figure 4.**
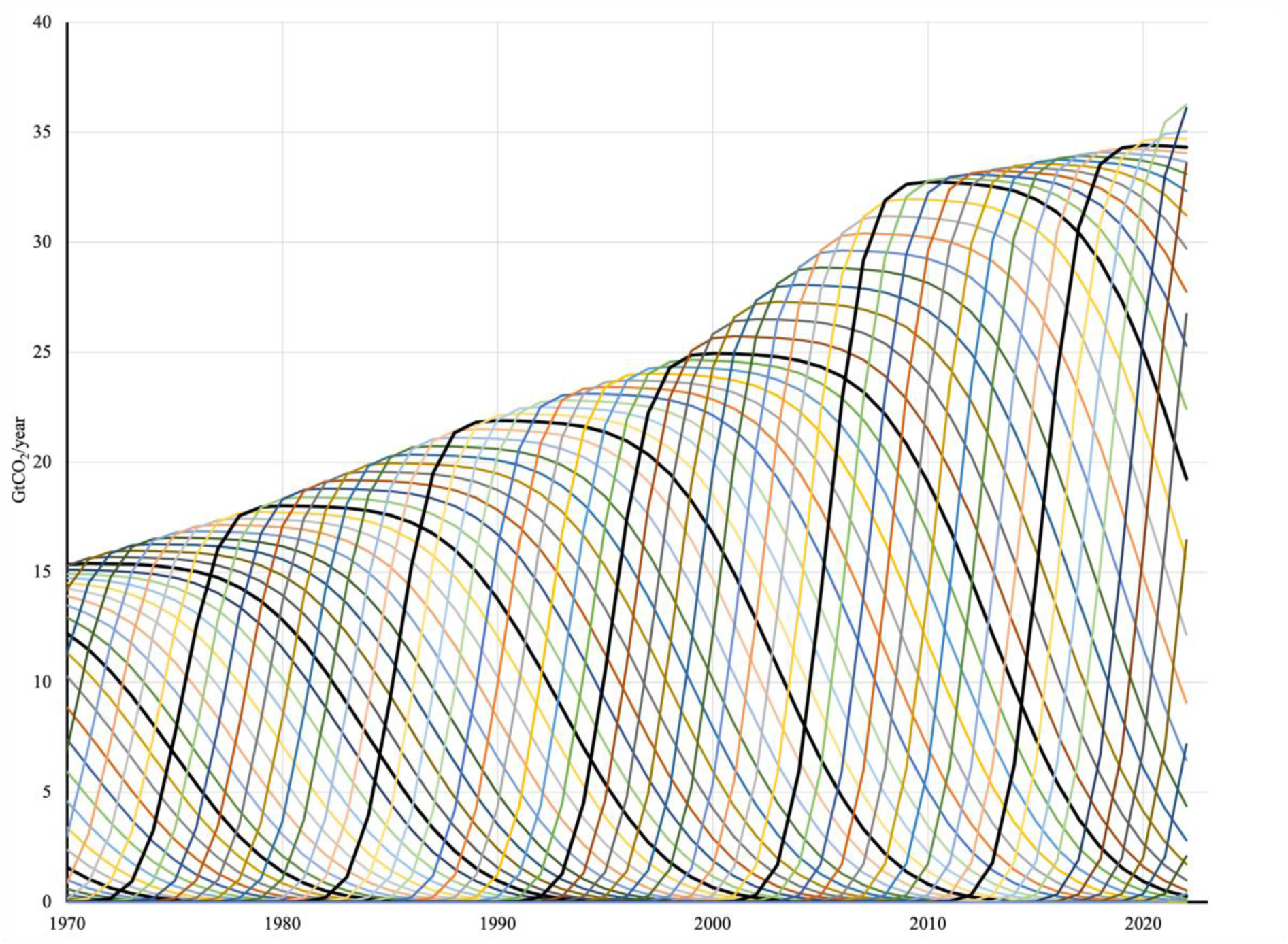
: Kinetics of carbon capture and storage by cultivated whole plants (CCSP) worldwide between 1970 and 2023 (GtCO_2_/year). The thick black lines correspond to the ten-year period.

### 9. Stocks from cultivated plants

The CSCPs of successive annual harvests shown in Figure 4 constitute the anthropogenic carbon stocks biofixed by cultivated plants present in the soil, buildings, stores and on shelves. To calculate these stocks accumulated in year n, we summed the elements of the row vector for year n in the matrix [C(t,n) + R(t,n)] from the column for year n minus 40, for which capture has not yet begun, to the column for year n. Figure 5 shows the evolution of these stocks over the half-century. The final downward hook could be related to the COVID- 19 pandemic.

**Figure 5.**
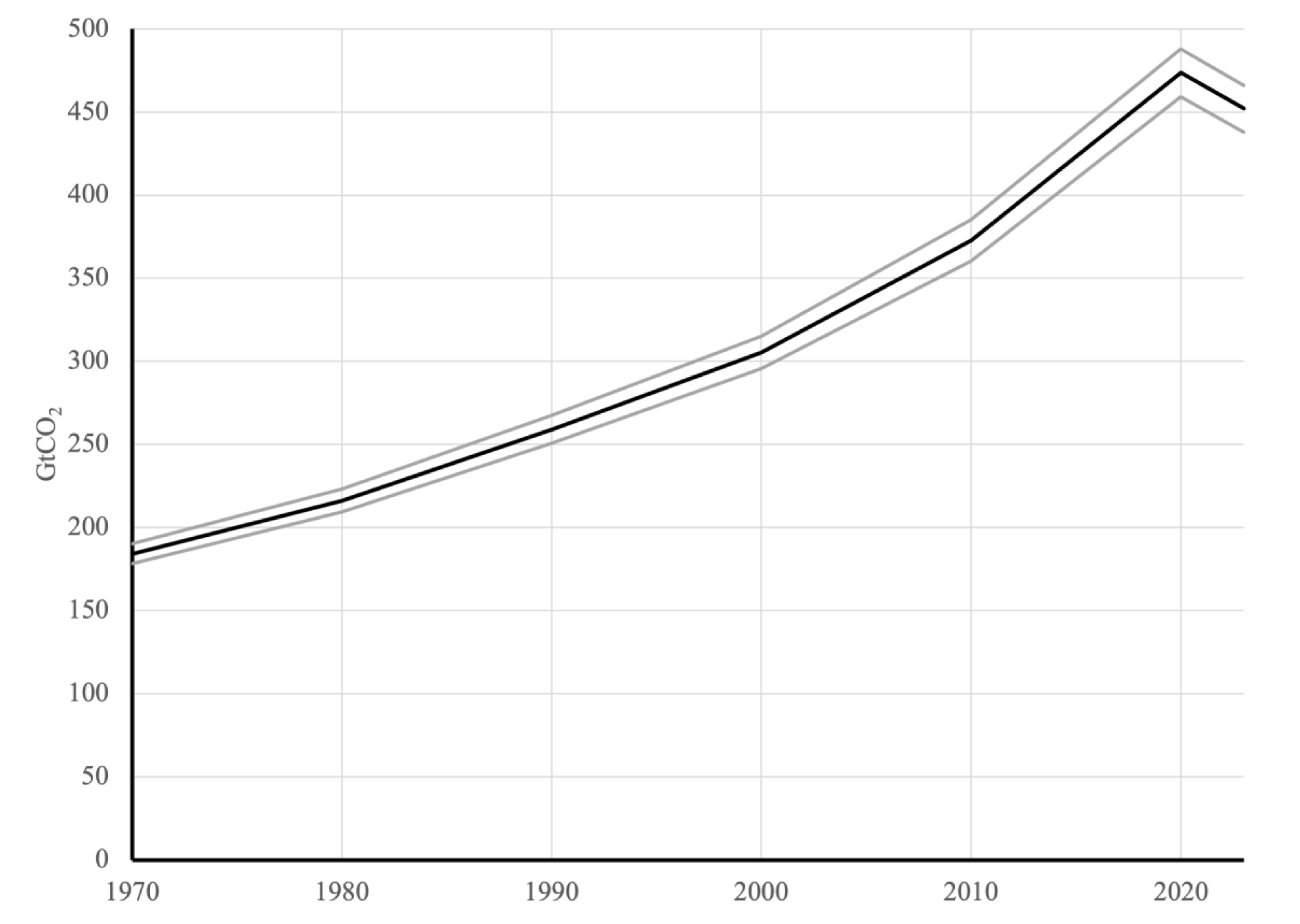
: Evolution over the half century of the cumulative net quantities of CO_2_ taken up by cultivated plants including restitutions. Average in black, plus and minus the standard deviation in grey.

In 1970, carbon stocks biofixed by cultivated plants had a net cumulated total of 184±6 GtCO_2_, or 7% of the atmospheric CO_2_ mass at the time. In 2023, these figures increased to 452±14 GtCO_2_ and 14%. Over the half-century, the carbon stock of cultures was multiplied by 2.5, as much as fossil emissions (2.5) and more than the human population (2.2).

The other components of the carbon budget identified in expression (2) are specified here. These are the emissions from fossil fuel combustion (EFOS) and the variation in mass of atmospheric CO_2_ (ACV).

### 10. Fossil fuel emissions

Figure 6 shows the distribution of CO_2_ emissions from fossil fuel combustion over the last half-century for selected countries and groups of countries, as provided by the UK-based Global Change Data Lab (GCDL). Global emissions reached 37.8 GtCO_2_/year in 2023, an increase despite a decline in 2020 during the health crisis. The main emitting countries are China (11.9 GtCO_2_/year), the United States (4.9 GtCO_2_/year), India (3.1 GtCO_2_/year), the European Union of 27 (2.5 GtCO_2_/year) including France (272.53 MtCO_2_/year). It seemed interesting to also look at a country with a low level of industrialization and seeking to escape poverty. We chose Kenya (21.5 MtCO_2_/year) because of its African position where countries with this profile are in the majority.

**Figure 6.**
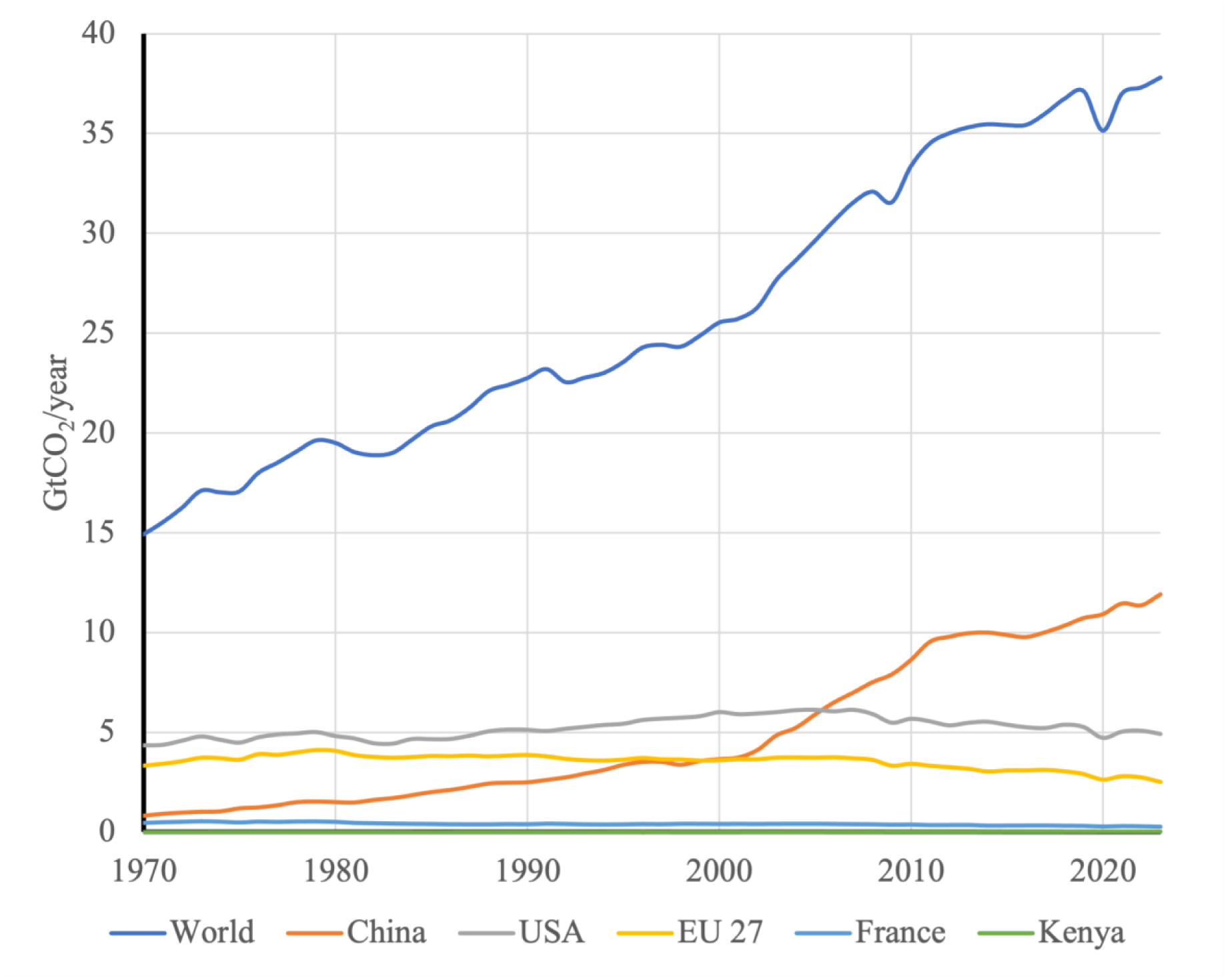
: Half-century variations in carbon dioxide emissions from fossil fuel combustion, in GtCO_2_/year (Source: Global Change Data Lab, n.d.).

#### World

While the share of the most industrialized countries, which are increasingly relying on low-carbon energy sources, has been declining since 2010, that of other countries, including China and India, which are actively equipping themselves with combustion-fired power plants, has been increasing since 2000. These emissions include those from transportation and other human activities, including agriculture, and other greenhouse gases (GHGs), which are grouped under the term “land-use change,” without CCSP.

To achieve carbon neutrality, most governments are developing CCS capacity, primarily through deep geological burial. As of July 31, 2023, the total capacity of global CCS by burying projects under development, construction, and operation was 361 MtCO_2_/year, or 1% of fossil fuel emissions. This modest capacity is up nearly 50% compared to 2022, according to the Global State of CCS report (Global CCS Institute, 2023). With carbon neutrality seemingly out of reach in the short term, a 55% emissions reduction milestone is planned for 2030.

CCS by burying involves a complex technological chain. In particular, CO_2_ capture before or after combustion has been the subject of research for several decades. Its technology involves amine-based solvents or membranes. Almost all solutions involve injecting the resulting CO_2_ into oil or gas wells. Geological storage sites are also former mines or diapirs. However, not all of these solutions allow for the recycling of CO_2_ after centuries of storage. It is an acid that reacts in the presence of water and combines with the geological substrate under the effect of temperature and pressure, contributing to its lithification.

Depending on the project, the costs of CCS by burying vary between €85 and €130 per tCO_2_/year, with CO_2_ extraction accounting for between 50 and 80%. The most active countries in this field are those with the most advanced development (United States, EU, United Kingdom, Japan), or those that produce oil (Saudi Arabia, Qatar, United Arab Emirates).

#### USA

As of September 2023, 15 CCSs by burying were in operation, realising a total of 22 MtCO_2_/year, or 0.4% of national emissions. The Congressional Budget Office (2023) allocated $5.3 billion from 2011 to 2023, primarily managed by the Department of Energy (DOE), for CCS research and development. There is also $3.4 billion from the American Recovery and Reinvestment Act in 2009, and $8.2 billion from the Infrastructure Investment and Jobs Act of 2021 for the period 2022-2026. In addition, companies that capture and store CO_2_ receive tax credits for each tCO_2_ sequestered. This represents a strong incentive for CCS, as US$1 billion in tax credits were requested between 2010 and 2019. A US$5 billion envelope from the federal budget is earmarked for these tax credits from 2023 to 2027. The Inflation Reduction Act concludes that it could increase CCS deployment in the United States from 200 to 250 MtCO_2_/year by 2030. The DOE Carbon Negative Shot aims to reduce the cost of carbon removal to $100 per tCO_2_/year. The EPA (Environment Protection Agency) is responsible for ensuring that storage is effective and sustainable. Agriculture and forestry are not identified in this context.

The incentive for CCS development was strong in the United States before President Donald Trump came to power. The CO_2_ emissions reduction policy implemented by the previous administration is being called into question. Instead, the government is encouraging manufacturers to increase their production and consumption of hydrocarbons.

#### European Union at 27 (EU)

According to the GCDL, the EU’s total fossil emissions have been on a downward trend since 1980. By 2022, they were down 31% compared to 1990, according to the European Environment Agency (2025). At the end of 2020, the EU adopted a new climate target for 2030, calling for a 55% reduction in its fossil emissions compared to 1990, a target of 2.13 GtCO_2_/year. The President of the European Commission stated in March 2023 that the EU should permanently store 300 MtCO_2_/year by 2050, which seems difficult to achieve without considering CCSP by plants. The latest target is “at least 50 MtCO_2_/year in 2030.”

The average price of carbon credits in Europe resulting from auctions in a market that generated €19 billion in 2020, rose from €37/tCO_2_ in February 2021 to €90/tCO_2_ in March 2023. The EU aims to set a carbon credit price high enough to encourage emitting industries to reduce their emissions. The revenues are distributed to Member States, which must use at least 50% of the proceeds for climate and energy-related measures, and 100% for aviation-related allowances.

#### France

During his speech on December 11, 2023, in Toulouse, the President of the French Republic identified CCS as a major challenge, with the goal of reducing industrial fossil fuel emissions by 10% by 2030. As 88.5 MtCO_2_/year are eligible for CCS, the stated objective is to implement CCS of 8.9 MtCO_2_/year. For example, a pilot project emitting 4,000 tCO_2_/year of gases from steel production has just been commissioned at the ArcelorMittal (n.d.) site in Dunkirk. The gases first circulate in two 20-meter-high columns through a chemical solvent that binds the CO_2_. The next step regenerates the solvent and recovers CO_2_ that is more than 90% purified. The purified gas is then compressed or liquefied for transport to a geological storage or utilization site. Onshore storage sites in France were identified in 2020 by ADEME (2020), which revealed a potential of 24 MtCO_2_/year, higher than the announced target. At that time, the contribution of forests and oceans to reducing emissions was mentioned, but not that of agriculture. The costs of CCS via onshore burial in France were estimated by ADEME at €100-150/tCO_2_ in 2020, significantly above European market prices.

#### Kenya

Kenya was chosen here to illustrate the situation of many countries seeking to escape poverty. According to the World Bank (n.d.), this country is “lower-middle-income, the only one in East Africa, the others being poorer.” With 54 million inhabitants, this country ranks 164th in terms of GDP per capita, at $2,099. Known for its safaris and tourism, its economy has suffered severely from the COVID crisis.

### 11. The atmosphere

The variation in atmospheric CO_2_ content (ACC) measured by Keeling C.D. (2001) at the Mauna Loa Observatory (MLO) on the Big Island of the Hawaiian archipelago since 1958 shows a clear upward trend of more than 2 ppm/year and exhibits annual oscillations with an average increase of 6 ppm from August to April (the boreal cold season) and an average decrease of 4 ppm from the warm April to August (the boreal season). Although located far from the coast and at high altitude, these records indicate a strong influence from the Northern Hemisphere continents, which emit the largest quantities of this CO_2_ and actively capture it through photosynthesis. Figure 7a depicts the kinetics of atmospheric CO_2_ stock. The smoothed CO_2_ flux to the atmosphere denoted ACV varies linearly as shown in Figure 7b.

**Figure 7.**
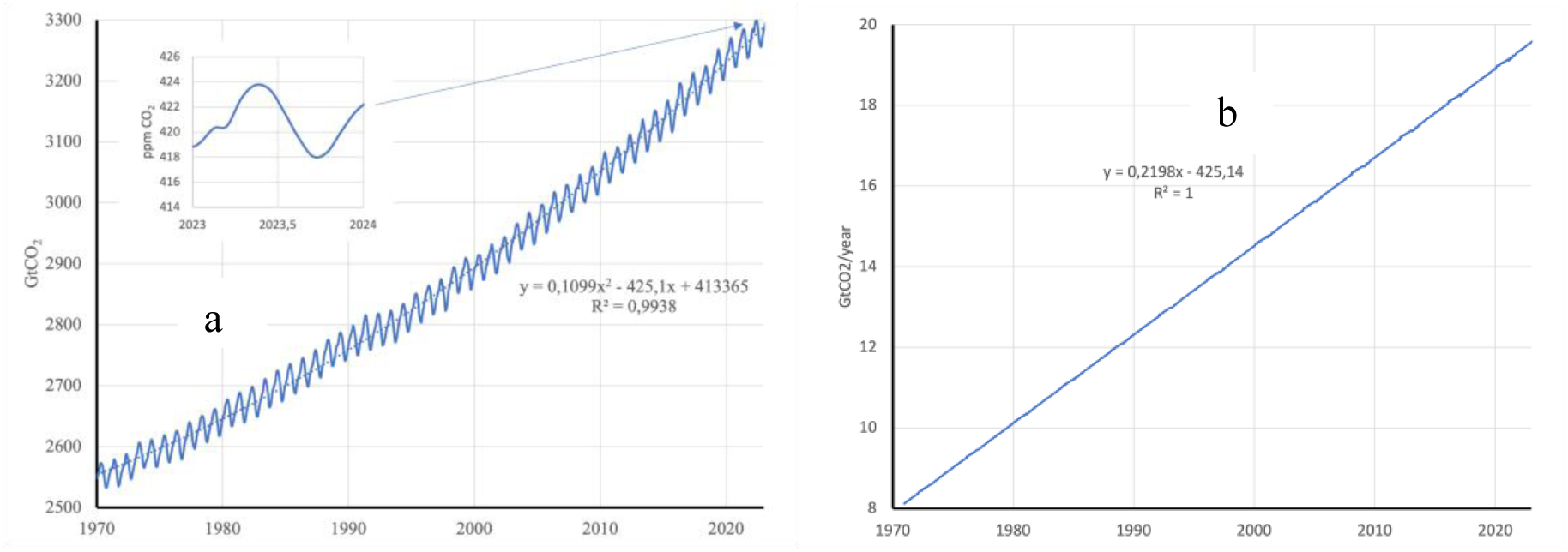
: Evolution of the mass of atmospheric CO_2_ in GtCO_2_ over the half-century on the left (a) with details of the year on the top 2023 (ppm), and its smoothed annual increase (ACV) on the right (b) in GtCO_2_/year, based on observations from MLO.

To determine the variation of atmospheric CO_2_ mass ACV, we multiplied the ACCs measured by the MLO by 7.84 GtCO_2_/ppm, then smoothed those values by a polynomial of order 2 (Figure 7a) and took the derivative (Figure 7b), i.e. the linear regression ACV = 0.2198 • n - 425.14 (R² = 0.99), where n is the date in years, which gives 19.52±0.25 GtCO_2_/year in 2023.

### 12. Carbon budget

The global budget according to formula (2) of CO_2_ exchanges in 2023 (Table 7) shows that plants exploited by humans are the planet’s main CO_2_ sink. The weighted average half-life DSC of this sink is 17.6 years and exceeds 40 years for forest products. According to our modelling, carbon capture by cultivated plants could offset a very significant—or even the majority—of fossil EFOS emissions, which calls for a reassessment of the respective roles of carbon sinks. The net plant balance –AW + EW of CCSPs was a sink of -31.0±1.9 GtCO_2_/year, offsetting 82% of fossil EFOS emissions.

**Table 7:**
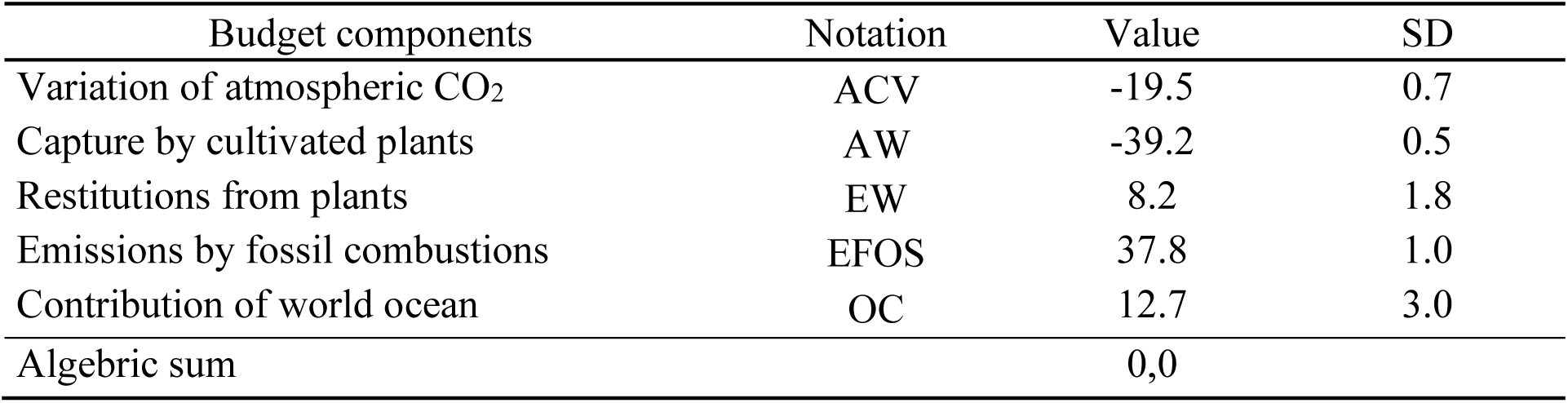
Elements of the budget of CO_2_ exchanges and their standard deviation (SD) in 2023 (GtCO_2_/year). Sources are positive and sinks negative.

| Budget components | Notation | Value | SD |
| --- | --- | --- | --- |
| Variation of atmospheric CO<sub>2</sub> | ACV | -19.5 | 0.7 |
| Capture by cultivated plants | AW | -39.2 | 0.5 |
| Restitutions from plants | EW | 8.2 | 1.8 |
| Emissions by fossil combustions | EFOS | 37.8 | 1.0 |
| Contribution of world ocean | OC | 12.7 | 3.0 |
| Algebraic sum |  | 0,0 |  |

The components of the global budget according to formula (2) vary as illustrated by Figure 8 during the last half century. With the exception of the restitution EW, these components are increasing in absolute value.

**Figure 8.**
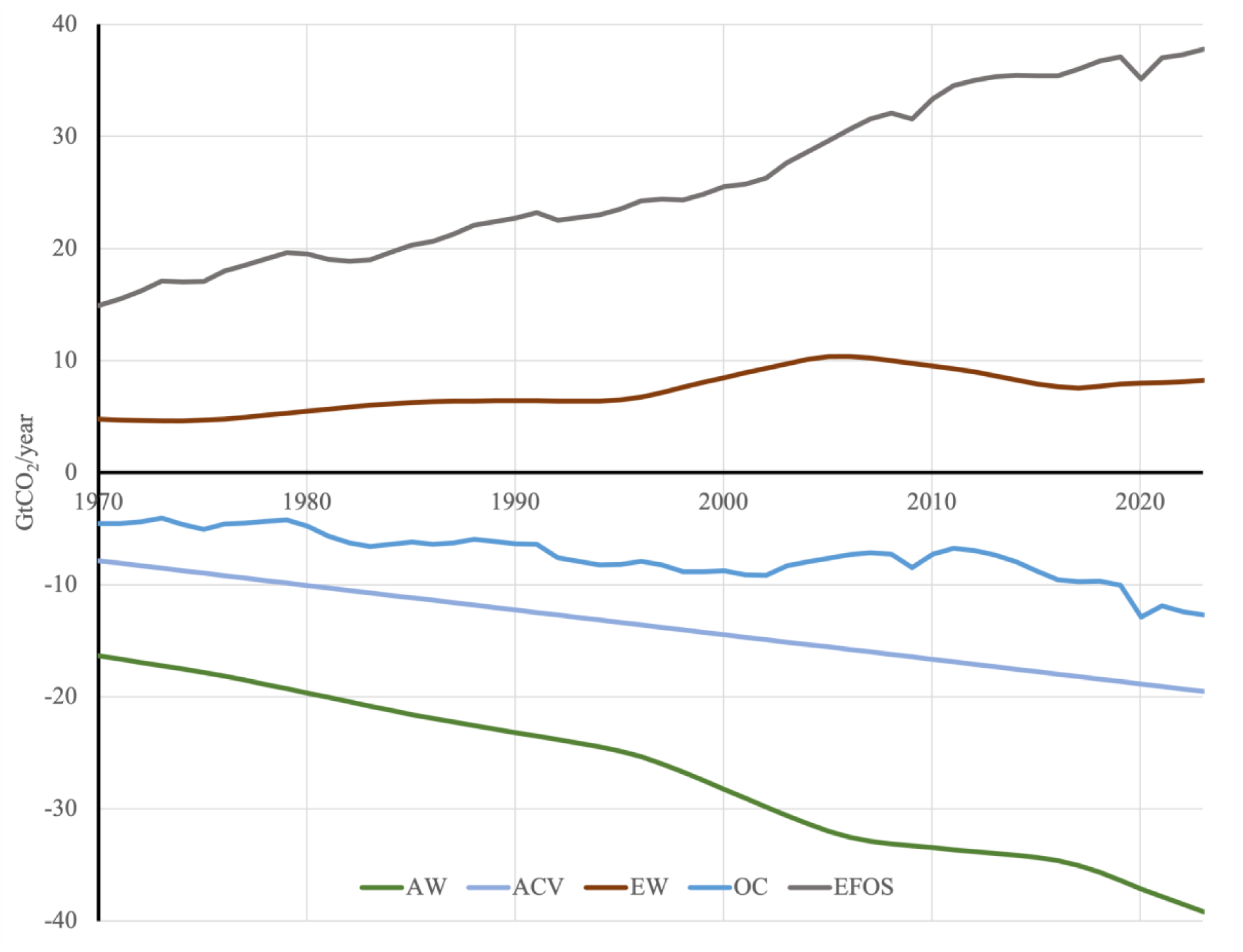
: Variations over the half-century of the five components of the CO_2_ exchange budget (AW plant captures; ACV atmospheric variation; EW restitutions from plants; OC ocean contribution; EFOS fossil emissions). Sources are positive and sinks negative. Their algebraic sum is zero.

On average over the half-century, the net balance – AW + EW of CCSPs was an anthropogenic sink of - 19.7±1.9 GtCO_2_/year, offsetting 75% of EFOS fossil emissions.

## Discussion

Absorption by cultivated plants (AW) and fossil hydrocarbon combustion emissions (EFOS) followed the evolution of the human population, which increased from 3.7 billion in 1970 to 8.1 billion in 2023. The three quantities more than doubled over the period while maintaining the same proportions.

This analysis highlights significant differences with the results presented in the literature. For the year 2023 alone, Figure 9 compares the components of the atmospheric CO_2_ budget resulting from our calculations with those proposed by Friedlingstein et al. (2023).

**Figure 9.**
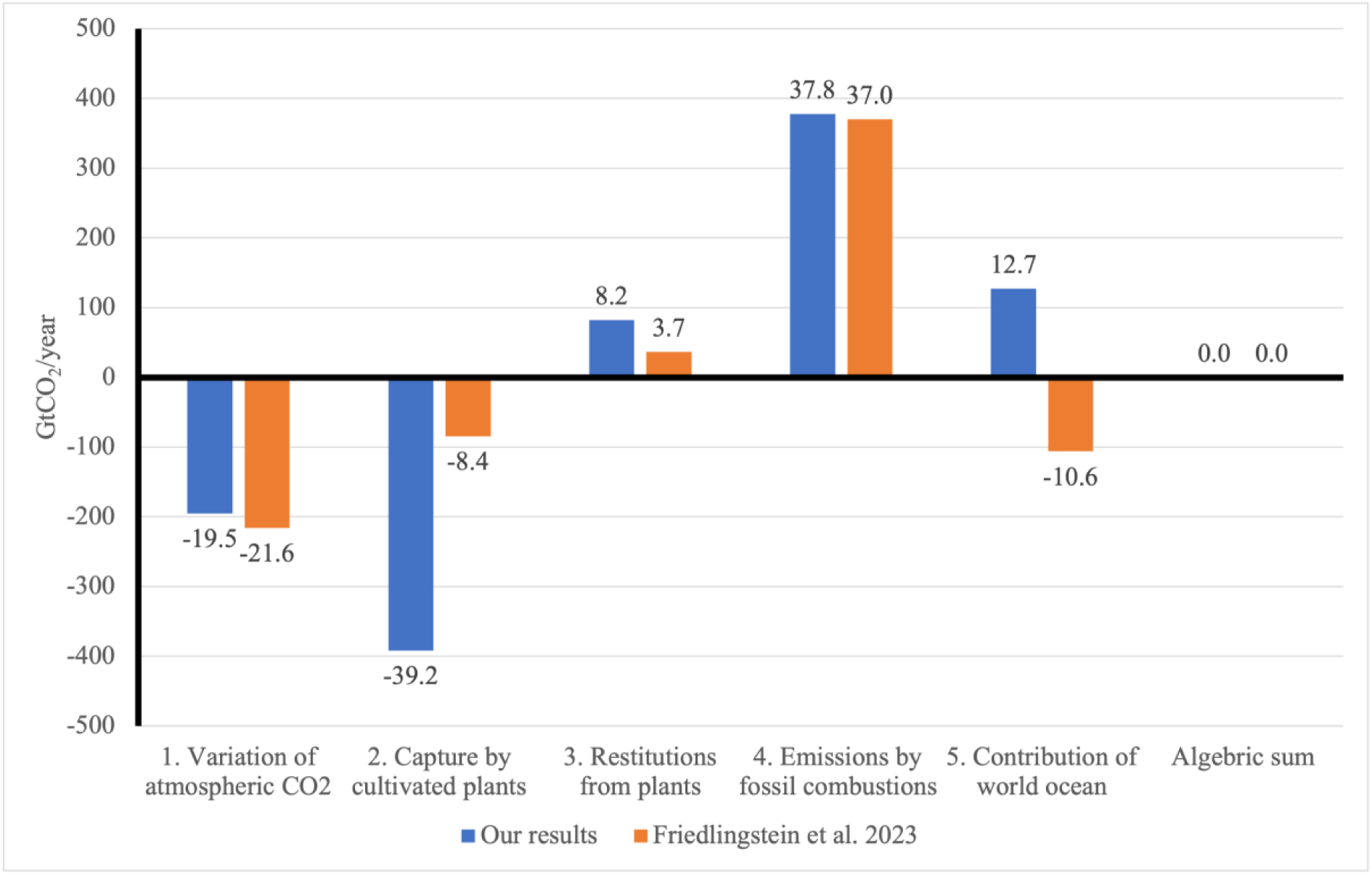
: CO_2_ sinks (negative) and sources (positive) calculated here and compared to those of the cited authors excluding annual crops, in billions of tons of CO_2_ per year for the world in 2023. The algebraic sum is zero in both cases.

In Figure 9, components 1 (ACV) and 4 (EFOS) obtained from public statistics, are similar. However, there are significant differences between components 2 (AW), 3 (EW) and 5 (OC). Our estimate of capture AW, which includes annual crops, is significantly higher (x5) than those excluding them. It was three times higher than that of Pan et al. (2024) estimated at 12.8 GtCO_2_/year during the 2010s. Let us recall here that our plant balance – AW + EW was a net sink of –31.0±1.9 GtCO_2_/year in 2023.

The capture by cultivated plants almost entirely offset alone total anthropogenic emissions (including non-fossil fuels) estimated at 41.46 GtCO_2_ by Ritchie & Roser (2024) in 2022, including 37.15 GtCO_2_ from energy and industrial combustion of fossil hydrocarbons (Global Carbon Budget). Jia et al. (2025) estimated the organic carbon sink constituted by global soils between 1992 and 2020 at 6.7±3.3 GtCO_2_/year. The figure put forward by these authors is of the same order as that of Friedlingstein et al. (2023).

Our restitution from plants 3 is twice that of the cited authors. Our contribution from the global ocean 5 is a source, whereas it is a sink for the cited authors. These differences result directly from the almost universal absence of annual plants in literature budgets, regardless of any other consideration. They are likely to change the way we view certain aspects of climate policies adopted to date.

### 1. Other approaches

Given the significance of the differences between our values and those in the literature, we verified their relevance using several approaches. We found only one publication in the literature describing the share of annual plants in global CCS (Wolf et al., 2015). Based on the same sources as us (FAO n.d.), their net primary production of harvested annual plants and forages required 28,2 GtCO_2_/year in 2011, compared to our 27,6 GtCO_2_/year for 2010, which is quite consistent. The orders of magnitude were three times higher than those of authors not taking annual plants into account (e.g. Le Quéré et al., 2012). However, Wolf et al. (2015) considered that the storage period did not exceed one year, with restitution by respiration immediately following harvest, which is consistent with the provision excluding annual plants. We consider that a half-life of carbon storage (DSC) of more than 10 years does not justify this exclusion and that annual plants have their place in carbon budgets.

Knowing the absorptions by cultivated plants AW and their DSC, we might expect the restitutions from plants EW to be close to the AW / DSC ratio, or 2.2 GtCO_2_/year in 2023. This figure is significantly lower than our EW value, probably due to the asymmetry of the kinetics around harvest.

We approximated the order of magnitude of the restitution by cultivated plants EW by considering global dioxygen consumption. According to Huang et al. (2018), it would be about 48 GtO_2_/year in 2023. The main contribution was due to the combustion of fossil hydrocarbons which consumed 38.2 GtO_2_/year in 2015, or 41.9 GtO_2_/year in 2023 modulated by the world population. The difference with the total global consumption of dioxygen, or 6.1 GtO_2_/year, is due to the mineralization of organic matter by respiration, fermentation and combustion. According to formula (1) taken in the direction of respiration from right to left, the CO_2_ production resulting from this consumption of oxygen would be 8.4 GtCO_2_/year, which is within the range of our EW assessment.

### 2. Methodological differences

Our capture by plants is higher by a factor of 3 to 5 than those in the literature and we are trying to understand the reason for such a difference. The first reason is the failure to consider annual plants on the pretext that their restitution would take place in the year following the harvest, which would cancel the capture. We have shown that this is not the case, the average half-life of annual plants weighted by their dry weight being 8.9 years. That of forage plants, most of them are annual, is 11.2 years. These durations are much longer than the year, taking away the meaning of the exclusion of annual plants which resulted in a strong reduction of the plant share. Also the net capture-restitution balances by plants are - 31.0±1.9 for us against -4.8 GtCO_2_/year for Friedlingstein et al. (2023).

There remain differences in the method of measuring quantities such as tree biomass and necromass. Our approach is based on the chemical composition of commercial products listed in FAO statistics. Most authors’ carbon stocks are constructed from field and satellite measurements per unit area and time, then extended by dynamic models to the global surfaces of the different plant covers thus characterized. It seems that this method faces problems of representativeness of the measurements characterizing each plant cover rather than the surfaces of these covers which are precisely defined thanks to satellite imagery.

Estimates of continental and oceanic absorptions and emissions lack precision, which encourages controversies (e.g. Luyssaert et al., 2008; Gundersen et al., 2021; Luyssaert et al., 2021; Zhong et al., 2024). In our case, the sampling is extended to all commercially available products from plants, to which should be added the proportion of unharvested plants, if significant.

### 3. Seasonal variations

The air mass surrounding the ACC measurement point at the MLO travels approximately two times around the planet between summer and winter, passing over most regions of the world. During this journey, it mixes at altitude and latitude and tend to reach equilibrium with surface water (Jiang et al., 2012). Figure 7 shows that ACC increases during the boreal cold season and decreases during the warm season. Several hypotheses have been put forward to explain these oscillations in ACC, which are out of phase with those of temperature. Some authors believe that they result from the high CO_2_ demand induced by photosynthesis during the boreal warm season, and increased emissions during the cold season in the northern hemisphere, which is the most industrialized and populated, thus favouring the influence of the continental biosphere. Other authors (e.g., Schindler & Bailey, 1993; Myneni et al., 1997) attribute these oscillations to biological activity, photosynthesis, and respiration. Some authors (e.g. Buermann et al., 2006) cite a North American sink in summer and a Eurasian source in winter.

We must also look to the ocean because oceanic primary production (OPP) is much greater than that of the continents. Aquatic life is primarily the work of ectothermic organisms whose metabolism is influenced by the temperature of the ambient medium. The restitution of CO_2_ to the atmosphere by heterotrophic mineralization occurs mainly during the warm season and could take precedence over physico-chemistry. In addition, the southern hemisphere has a much larger maritime surface than the northern hemisphere. The influence of the former could be manifested at the MLO (19.53° north latitude) with reversed seasons. The oceanic regions of the two hemispheres which are the site of the largest average CO_2_ exchanges are located as a source (water to air) around the equator and as a sink (air to water) around the 40th parallel (DeVries et al., 2023), which further reinforces this asymmetry in favor of the southern hemisphere.

### 4. The Ocean

According to our calculations, the ocean would be an increasingly important source of CO_2_ (Figure 8). It would release 12.7±3.0 GtCO_2_ in 2023 (Figure 9) and would account for the atmospheric increase with fossil emissions and plant restitutions. The role of a sink of 10.3±0.4 GtCO_2_/year attributed to the ocean (e.g., Fay et al., 2023) by authors who excluded annual plants would thus be called into question in both sign and magnitude. By choosing the notation Socean for “ocean sink”, these authors prohibit themselves from conceiving the ocean other way than a sink (e.g., Rödenbeck et al., 2015). This conclusion appeared to be corroborated by numerous offshore measurements of CO_2_ fugacity and total inorganic carbon, which tend to show that the ocean absorbs a quarter of anthropogenic emissions (NOAA n.d. - SOCAT). Thus, offshore measurements may be insufficient to describe mass exchanges due to gaps in space and time, as well as limitations related to the accuracy of sensors and interpolation models, which fuels controversy (e.g., McGillis et al., 2004; Crisp et al., 2022).

The ocean is believed to be degassing, probably due to warming, which reduces gas solubility. According to ice cores, CO_2_ peaks follow temperature peaks 600 to 1,000 years later (e.g. Petit et al., 1999; Fischer et al., 1999; Caillon et al., 2003; Richet, 2021). This would be the time it takes for the global ocean to find a new thermal equilibrium with the atmosphere through mixing in all three dimensions. The ocean’s delayed response to climate perturbations induce to look for the warming that caused the current outgassing well before the recent increase in global temperatures. The so-called “Medieval Climate Optimum,” which occurred from the 10th to the 13th century, could be the cause of this outgassing, which was briefly interrupted or slowed by the “Dalton Minimum,” or “Little Ice Age” (1790-1830).

The OPP provided by algae is the basis of the aquatic food chain exploited by marine fisheries, which landed 81 million tonnes of products in 2022, stable since 1980. Aquaculture, increasing by more than 10% per year, produced 185.4 million tonnes of animals and 37.8 million tonnes of macroalgae in 2022 (FAO, 2024). OPP has been the subject of numerous studies, mainly using satellites measuring chlorophyll. Since 1970, most authors agree to consider OPP as stable and equal to 51 GtC/year (e.g. Carr et al., 2006; Johnson K. S. & Bif M. B., 2021), with only events modifying deep water upwellings (e.g. El Niño) disrupting the annual rhythm. This is 4.8 times our estimate of uptake by cultivated plants AW, even though the ocean surface area is 2.4 times larger than that of the continents. Relative to surface area, ocean capture exceeds that of the continents by a factor of two. This demonstrates the extent to which aquatic life plays a key role in the global terrestrial ecosystem, consuming carbon and producing the oxygen and organic matter essential for animal life.

Vertical profiles of inorganic carbon have been established in all ocean regions and in all seasons. They exhibit different concavities depending on whether the ocean is a source or a sink. In the former case, the surface is less rich than the depth, and in the latter case, it is the opposite. Source regions are mainly located on the western shores of continents where deep waters upwell. Sink regions are located at high latitudes where colder and denser surface waters sink. According to McKinley et al. (2023), oceanic CO_2_ is thought to be of deep origin. Vertical exchanges of inorganic and particulate carbon between the depths and the surface of the ocean ensure its renewal to establish equilibrium with the troposphere. Part of the OPP and the animal biomass that feeds on it passes into the deep ocean in particulate form by gravity and transport. On the scale of an annual cycle, Falkowski et al. (1998) estimated the primary production sinking to the bottom at 16 GtC/year, or one third of the OPP. For their part, Levi et al. (2013) estimated the exchanges from the depth to the surface at 275 GtC/year and, in the opposite direction, at 265 GtC/year, which leaves a gain of 11 GtC/year for the surface ocean. Thus, the balance between the surface and deep ocean gives rise to various assessments which show the difficulty of the exercise.

### 5. Seasonal and Glacial Cycles

During glaciations, ice accumulated at the high and mid-latitudes of the continents. Several thousand meters of thickness thus shaped their relief, while sea level was more than 100 m below current levels. The increase in the ACC following glaciations is a consequence of the degassing of CO_2_ from the ocean, which is a slow process involving the deep oceans. These cycles differ from the seasonal cycles of the ACC, whose oscillations are in opposition to those of temperature.

According to ice cores, the atmosphere would lose 100 ppm, or 212 GtC, while the temperature would drop by about 8 C°C on average during glaciations (Kobashi et al., 2013). This carbon mass would pass into the ocean, which would release it during postglacial warming. These exchanges represent a tiny fraction (0.5%) of the oceanic carbon stocks estimated at 40 TtC (Bopp et al., 2019). Since the mid-19th century, the atmosphere has been enriched by 300 GtC. Thus, during this short period, the losses from the last glaciation were largely offset by the two main sources: fossil emissions and the ocean.

Seasonal cycles are due to the obliquity of the Earth’s axis relative to the ecliptic plane, while glaciations are primarily the effect of changes in Earth’s orbit around the sun. The seasonal temperature amplitude is of the same order as that of glacial cycles. Consequently, CO_2_ solubility variations for the two cycles should be similar. However, the amplitude of the ACC is 3.1 ppm for a seasonal cycle and 100 ppm for a glacial cycle. What differs between these cycles is the duration of the disturbances: a few months for the first versus a few millennia (ky) for the second. In the case of the seasonal cycle, the physical equilibrium does not have time to stabilize before the next cycle and the influence of the biosphere is predominant.

Whereas for glacial cycles, thermal exchanges take place over a sufficient duration for this equilibrium to be reached, with physics and chemistry prevailing over the biosphere.

### 6. Surface capture yields

The global capture figures from our calculations (Table 6) are understated because they do not include the share of non-commercially exploited plant covers. These cover types include forests (tropical, temperate, boreal), tundra, Mediterranean scrub, tropical savannahs and prairies, temperate prairies, and deserts. According to Saugier et al. (2001), surface capture would range from 3.7 (deserts) to 43.1 (tropical forests) tCO_2_/ha/year. The share of crops was only credited with 10.1 tCO_2_/ha/year by these authors.

The share of forests was greatly overestimated, and that of crops was significantly underestimated. Indeed, we estimate the average yield per unit area of CCSP, weighted by the dry weight of cultivated whole plants, at 19.4±0.7 tCO_2_/ha/year worldwide in 2023. Sugar crops (cane and beets) and palm oil lead the way with two to three times the average, followed by vegetable crops (onions, carrots, turnips, beans, eggplant, etc.), whose yields are above average.

Unexploited continental vegetation is stable and plays a marginal role in capturing atmospheric carbon. Should unexploited old-growth forests still capture carbon, they would do so at levels far below those of most crops and forages. For example, current logging regulations in Brazil adopt a standard post-harvest recovery rate corresponding to a surface capture yield of 0.9 tCO_2_/ha/year (Vidal et al., 2020), 20 times less than crops. The average capture of French exploited forests is 3.3 tCO_2_/ha/year (IGN, 2013), 6 times less than crops.

Forest logging and fires do not result in a loss of capacity to capture and store CO_2_. The soil stock remains in place, while spontaneous or replanted regrowth continues to capture CO_2_ before further logging 25 to 35 years later. If the forest is replaced by pastures or crops, the space it frees up captures CO_2_ at levels more than 10 times higher.

### 7. Climate

Carbon dioxide (CO_2_) is the natural constituent of the atmosphere to which we owe the main forms of life. Its combination with water during photosynthesis produces the plant biomass and oxygen necessary for heterotrophic life. The ACC has been stable at 260±15 ppm since the end of the last glaciation 13 ky ago and began to increase sharply a century and a half ago to exceed 425 ppm. This growth of 2.2±0.1 ppm/year on average is assumed to be linked to human activities burning fossil hydrocarbons, natural gas, oil, coal, and to deliberate or accidental fires. However, we have seen that this fossil combustion is offset by plants, which calls into question the responsibility for these emissions and the resulting climate policy.

#### 7.1 Solar activity

Milankovitch (1913) has shown that the cycles of Quaternary glaciations result from variations in the precession of the equinoxes (conical rotation of the Earth’s axis of rotation around an axis perpendicular to the ecliptic plane (main period of 26 ky), the inclination of the Earth’s axis of rotation relative to the ecliptic plane (obliquity) from 22.1 to 24.5° (main period of 41 ky), the eccentricity of the Earth’s orbit (main period 100 ky), and solar radiation. Carbon 14 has made it possible to trace the activity (especially magnetic) of the Sun over the last centuries and millennia and to compare it with the climatic variations recorded by historians.

Measured for almost 3 centuries by the number of spots on the visible surface, the Sun’s activity has an average periodicity of 11.2 years. We are almost at the maximum of the cycle number 25, and a decrease in this activity is to be expected. In its 5th report, the IPCC reduces the sun’s effect solely to variations in its brightness. It is true that, during these solar cycles, the intensity of the radiation impacting the Earth’s periphery varies slightly, between 1360 and 1362 W/m^2^. However, other parameters should not be neglected, notably the movements of the center of the sun around the center of gravity of the solar system, which varies the Earth-Sun distance, the modulation of cloud cover by solar radiation, etc.

The question of a new ice age beginning before 2000 was discussed in the 1970s with the observation of a marked cooling between 1940 and 1950 and then between 1970 and 1980. Short-term forecasts cannot be based on Milankovitch’s theory, which does not provide any clear trend on a century-long scale. However, some climatologists suggest the preponderance of solar activity on climate over the past few millennia. They note that climate change and solar activity, measured by radiative energy impacting the Earth’s surface, are strongly correlated, and that the influence of CO_2_ on climate is secondary. Since solar activity has increased continuously since the beginning of the industrial era, some authors believe there will be a medium-term trend reversal towards a minimum like that of Dalton (”Little Ice Age”, 1790-1830).

#### 7.2 The Earth’s Past

Ice cores drilled at the Russian Vostok Station in East Antarctica provided the first climate record covering the last 423 ka (Petit et al., 1999). They indicate a glaciation periodicity of 80 to 110 ka and a strong correlation between ACCs and isotopic temperatures indicated by ice cores. During this recent history of the Earth, the lowest ACCs are observed during glacial periods and the highest during interglacial periods. However, this correlation does not clarify the cause and effect between ACC and temperature.

The theory that temperature drives ACC, and not the other way around, is validated by other considerations from the analysis of glacial archives (Richet, 2021): i) Temperature peaks last less long than those of the ACC, which indicates the driving of the ACC by temperature, a disturbance always lasting less or as long as its effects. ii) Two closely spaced temperature peaks correspond to a single ACC peak, which would not be possible if the first were induced by the second.

The ACCs were much higher in the Earth’s past, including during glacial episodes (Carboniferous - Permian). Thus, they reached 7,000 ppm (0.7%) in the Cambrian (540 million years ago - My) and still 1,800 ppm (0.18%) in the Jurassic (200 My), i.e. between 4.3 and 17 times higher than today. The temperature of the troposphere was significantly higher than today due to a greater mass linked in particular to these high levels of CO_2_, the heaviest gas of the atmosphere. Life has developed since the Precambrian without being interrupted by high temperatures.

#### 7.3 The Greenhouse Effect

The recent increase in the ACC is presented as contributing to global warming by preventing excess heat brought by the solar flux from being re-emitted into space, causing it to accumulate in the troposphere. Most models are based on the greenhouse effect theory conceived by Joseph Fourier in 1824 (Dufresne, 2006) and formulated by Svante Arrhenius in 1896 (Acot, 2008), according to which the temperature variation is proportional to the variation in the logarithm of the ACC. Constantly evolving, these models poorly simulate average annual variations in surface temperature over several decades. Most overestimate the warming trend observed in the tropical troposphere over the last 30 years and tend to underestimate the long-term cooling trend (Flato et al., 2013).

Other theories claim to explain the radiative dynamics of the atmosphere and its thermal behaviour. Miskolczi (2007) presents a theory of the greenhouse effect based on the virial theorem linking the kinetic and potential energies of several interacting bodies. He concludes, on the one hand, that the influence of greenhouse gases on global warming is overestimated by those who claim to be responsible for it, and on the other hand, that the increase in the contribution of one atmospheric component to the greenhouse effect is offset by the decrease in that of another component. In other words, the greenhouse effect would be permanently saturated.

By reasoning by the absurd, if we cross the observation that the ocean degasses CO_2_ when the temperature increases with the putative principle of a climate influenced mainly by the greenhouse effect of CO_2_, the increase in the ACC by oceanic degassing with warming would be a self-sustaining process since it would generate by greenhouse effect a new warming which would cause a new degassing of the ocean, etc. This divergent process, which would end with the boiling of the oceans, should leave traces in the past of the Earth. However, no sign of this runaway temperature has been observed despite ACCs much higher than today.

### 8. Carbon Capture and Storage (CCS)

According to the Global CCS Institute (2023), when all global landfill CCS projects are operational, their capacity will be 100 MtCO_2_/year, 440 times less than human-exploited plants, which constitute by far the largest CCS device on the planet.

All of the European Union’s emissions (2.51 GtCO_2_/year) were offset by its cereal production alone in 2023 (-3.5 GtCO_2_/year). The EU considers it a priority to find adequate financial rewards to promote carbon removal in the various sectors of agriculture, forestry, and industry, but has still not recognized CCS by plants. Several countries are aware of the important role of plant crops, but are nevertheless raising the question of certification, on a scientific basis accepted throughout the EU, of CO_2_ capture by plants and the duration of storage.

France is highly decarbonized, with most of its electricity being nuclear generated. Without any incentive or change in practices, French agriculture captured 174 MtCO_2_/year for 17.6 years, 64% of total national emissions in 2023, more than six times the CCS target for 2030, and twice the industrial emissions eligible for CCS.

The agricultural sector is criticized for its emissions of CO_2_, methane (CH_4_), and nitrous oxide (N_2_O). According to Maréchal (2022), for example, the sector is responsible for 23% of global anthropogenic greenhouse gas emissions, or 12 GtCO_2_ equivalent/year. These authors do not consider continental photosynthesis, whose net capture-restitution balance is largely favourable to the rural world, with 31.0 GtCO_2_/year removed from the atmosphere, offsetting 84% of fossil fuel emissions in 2023.

The failure to consider agriculture and forestry in CO_2_ emissions offsetting would be a persistent injustice if there were not some signs of recognition underway, marking a significant shift toward redressing this oversight, which harms the rural world. Thus, since 2020, the French government has identified a so-called voluntary carbon offset market, which provides that “…*farmers can put the carbon credits they have generated…on the voluntary carbon offset market*.”

At the France Grandes Cultures congress held in February 2023, APAD (Association for the Promotion of Sustainable Agriculture) announced a minimum price of €50 per tCO₂ stored (carbon credit) paid to the farmer. The average for large-scale crops is credited with capturing and storing around 1.5 tCO₂/ha/year, which is much lower than our estimate of around 19 tCO₂/ha/year for grain maize and 18 for wheat. Soil Capital estimates that €33 per tCO₂ will be paid to the farmer in 2022, which represents an average of €5,000 per farm per year. Based on an average of €100 per tCO_2_, CCSP remuneration would generate €23 billion of additional income, adding to the €57 billion in products sold in 2021, representing an increase in French farmers’ income of around 40%.

Kenya has a significant surplus and offsets its fossil fuel emissions more than three times over with plants. Like most countries seeking to escape poverty, it could sell its surplus to emitters through the global carbon market. For example, in the case of Kenya and a carbon credit of €100/tCO_2_, this could generate €9 billion per year for local farmers.

The consideration of CCSP is intended to extend to all plant products placed on the market. Storage duration varies from a few months (fruits and vegetables) to several centuries (construction timber). The remuneration of CCSP by plants would represent a significant source of income for farmers, whose mission would now be, in addition to providing essential foodstuffs for humanity, to avoid or offset CO_2_ emissions to limit climate change for which this gas would be responsible.

It should be noted here that there are other plant resources, cultivated or not, that could constitute CCSP. Thus, cultivated macroalgae captured 8.0 MtCO_2_/year worldwide in 2023. Sargassum, which naturally invades the equatorial Atlantic, could be harvested at sea and then transformed into biofuel and fertilizer, which would reduce the use of fossil fuels and, simultaneously, the disastrous strandings on the eastern coasts of Central America and the Caribbean islands. According to Marx et al. (2021), the processing of this sargassum could produce 8,500 barrels of crude oil per day for a CCSP of 1.9 MtCO_2_/year.

### 9. Carbon dioxide scarcity

Mainly as a result of human population growth, agricultural land per capita (ha/c) has halved during the last half century, from 1.2 to 0.6 ha/c. Despite this, per capita production has increased from 0.51 to 0.60 t/c/year, preventing food shortages. Plant production is experiencing a boom, as evidenced by leaf area measurements and the greening of the Earth (NASA, 2016). The increase in ACC from 325 to 420 ppm is the main cause of the improvement in overall agricultural production yields per unit area (Haverd et al., 2020), which more than doubled from 0.5 to 1.3 t/ha/year, according to 2019 FAO statistics (n.d.). This could also explain why the net cumulated carbon stock biofixed by cultivated plants has more than doubled, passing from 184 to 452 GtCO_2_ removed from atmosphere.

Carbon dioxide dissolves in seawater with which it combines to form bicarbonate and carbonate partially insoluble ions. Carbon is distributed in the deep oceans in dissolved form by density drifts and in particulate form by gravity and drifts. Insoluble carbonates accumulate at the bottom along with organic matter to constitute the planet’s main carbon stock in the mineral form of limestone rocks (1.4 105 TtC according to Sorokhtin et al., 2007) and in the organic form of fossil hydrocarbons (oil, coal, bitumen, and gas – approximately 150 TtC). The main coal reservoir was formed with the development of forests at the beginning of the Carboniferous period (-359 to -299 My) when the ACC was three to four times higher than in the present day.

Air-water exchange of CO_2_ is a powerful regulatory process of the ACC, including for photosynthesis, since this gas is admitted into the plant cell in dissolved form before being polymerized. Given that the ocean contains approximately 50 times more carbon than the atmosphere, the ACC will be strongly affected by temperature variations when these have time to express. The oceans will tend to absorb atmospheric CO_2_ in cold regions and periods and release it back into the atmosphere through outgassing in warm regions and periods.

The Earth’s early atmosphere was predominantly composed of CO_2_. The ACC has decreased by a factor of 2,000 since the Earth’s origin 4.5 billion years ago (Gy), and this decline continues inexorably at a rate of approximately 13 ppm per million years. It still reached 7,000 ppm in the Cambrian period (-540 My), then 1,800 ppm in the Jurassic period (-200 My). No catastrophic warming interrupted the development of life that appeared 3 Gy ago. The fact that glaciations sometimes coincided with high CO_2_ concentration levels (Carboniferous-Permian) is another indication of the absence of control of temperature by CO_2_.

This slow decrease in ACC by lithification will result in a cessation of life on Earth, if nothing interrupts it beforehand. Indeed, photosynthesis in plants with C3 photosystems (wheat, rice, barley, beans, cassava, soybeans, etc.) stops below 100 ppm and the yield of other plants (corn, sorghum, sugarcane, etc) decreases sharply. The CO_2_ breakdown was already approached when the ACC dropped to 150 ppm in the Carboniferous-Permian (-290 My) and then in the Pleistocene (-2.6 My to -11,700 years ago). The current rise in ACC is a respite from this slow decrease by lithification. If the anthropogenic origin is proven, it would result from the recycling of carbon trapped in the Earth’s crust, a kind of reverse lithification a million times faster. The decrease of ACC could also be partly offset by degassing of the Earth’s crust and volcanic activity (Burton et al., 2013). In any case, to prolong the availability of this precious gas, it is advisable to prefer CCSP to CCS by burial which contributes to its lithification.

## Conclusion

The contribution of cultivated plants to the biofixation of atmospheric CO_2_ is approached with a margin of uncertainty relating to both retention times and stored quantities. This is a delicate exercise due to the diversity of elements to be gathered, water content, storage times, quantities, which we have selected among the most relevant from the literature. When doubts arose, hypotheses minimizing both values were retained, which gives them a minor and conservative character. Subject to the precision and completeness of these elements and the quality of the statistics on which they are based, the CCSP by commercial products placed on the market in 2023 presented a half-life of more than 10 years, which makes it a component to be considered in carbon budgets. The same is true a fortiori when we include the non-commercial parts of whole plants which remain on site and enrich the soils with carbon and which had a half-life of 17.6 years. They capture twice as much carbon dioxide and retain this stock almost twice as long as the commercial products they carried.

The kinetics of carbon capture and restitution over the half-century were simulated using a probabilistic approach, which notably enabled the matrix calculation of intermediate capture and restitution through biomass mineralization. The total quantities of carbon captured and stored during these rural productions were five times greater than those reported in the literature. The contribution of annual crops to the CCSP was found to be 77% with an 8.9 years half-life. Among them, cereals alone captured 60% of annual crops and 21% of all plants over a half-life of 10.4 years. The remainder is due to perennial crops (fruit trees, oil palms, etc), the only ones considered by the authors cited in their carbon budgets.

These figures require further refinement and completion, precisions on their evolution over time and space, but they do depict major trends. They reveal the exceptional nature of the contributions of agriculture and forestry, which are not yet fully recognized. These activities mobilize atmospheric CO_2_ at levels that make them by far the main CCS system in the hands of humans, without humans having to modify their practices to achieve this result. Moreover, they allow the return to the atmosphere of this gas, which is crucial for the planetary ecosystem, unlike geological CCS, which contributes to its lithification.

According to our simulation, anthropogenic fossil emissions would not be responsible for the increase in the atmospheric carbon content since they were entirely bioremedied by the capture of cultivated plants. This calls into question their influence on climate change, and the policies for their reduction. In addition to fossil emissions and those by restitution from plants, the other source responsible for the enrichment of the atmosphere would be the ocean. Indeed, contrary to most assessments, our results suggest that the oceans may not play the commonly attributed role of carbon sink but constitute a source providing more than half of this increase.

Based on their significant carbon capture and storage durations, agriculture and forestry should be reevaluated by international organizations and governments. These ancestral activities, which feed, warm, cheer, and clothe humans, use photosynthesis, to which we owe our lives, and which remains the simplest and most efficient means of capturing and storing atmospheric CO_2_. They should be rewarded for their carbon credits as part of the development of CCS, which would supplement their income so they can continue to operate, develop, and transfer their activities.

The convention that carbon captured by annual plants would be cancelled out by its release in the year following harvest favours long-term stocks and neglects the variation of these stocks in stores, on shelves, and in the soil. By reducing the contribution of carbon fluxes from rural origin, the exclusion of annual crops from carbon budgets could lead to energy and climate policies disconnected from agronomic realities, and encounter incomprehension in the agricultural world and developing countries.

## Abbreviations

COP: conference of the parties
DOE: Department of Energy
EU: European Union at 27
GCDL: Global Change Data Lab
GDP: Gross Domestic Product
IPCC: Intergovernmental Panel on Climate Change
MLO: Mauna Lao observatory
OECD: Organisation for Economic Co-operation and Development
WMO: World Meteorological Organization
UN: United Nations
UNEP: UN Environment Programme

## Notations

AW: absorption by cultivated plants
Co: oceanic contribution
CP: photosynthetic capture
CCS: carbon capture and storage
CCSP: carbon capture and storage by plants
EFOS: emissions from fossil fuel combustion
EW: emissions coming from plants
DSC: half-life of carbon storage
OPP: ocean primary production
RM: restitution by mineralization
ACC: atmospheric CO_2_ content
ACV: variation in mass of atmospheric CO_2_

## Units

ppm: parts per million. 1 ppm of CO_2_ = 7.84 GtCO_2_
M: mega (10^6^)
MtCO_2_: million tons of carbon dioxide
MtC: million tons of carbon
G: giga (10^9^)
GtCO_2_: billion tons of carbon dioxide
GtC: billion tons of carbon
GtO_2:_: billion tons of dioxygene
Gy: billion years
c: inhabitant
ha: hectare (100 m * 100 m = 10,000 m^2^)
ky: kilo years - 1000 years
My: million years
T: tera (10^12^)
TgC: tera (10^12^) grams of carbon = MtC
P: peta (10^15^)
PgC: peta (10^15^) grams of carbon = GtC
W/m^2^: Watt per square meter

## Acknowledgments

We are particularly grateful to all the data collectors at FAO, GCL, NASA, BP, etc., without whom this analysis would not have been possible. We thank the many people involved in collecting and making the global data available and recognize the considerable resources in personnel and time devoted to collecting this invaluable data. We also thank the engineers who wrote the programs to process this data, put it online, and make it easily accessible to users.

